# Attachment site, not linker length, bounds tethered base editor windows

**DOI:** 10.64898/2026.09.16.751980

**Authors:** Anees Ahmed Mahaboob Ali, Everette Jacob Remington Nelson

## Abstract

Base editors act on the DNA strand displaced within an R-loop, and the set of positions they convert, the activity window, has been engineered for a decade on the assumption that linker length and attachment geometry determine it. We built a geometric model that predicts where a tethered deaminase acts from linker statistics, steric exclusion, and R-loop geometry alone; fitted three parameters to two previously reported profiles, held them fixed, and scored predictions against 50 architectures from seven studies, with each substrate coordinate withheld. Varying the contour length by a factor of 16 does not shift the predicted window at all, whereas changing the attachment site does: across 185 buildable single-linker designs at thirteen attachment sites, the predicted peak never leaves protospacer positions 3 to 12, positions 1, 2 and 13 to 20 are reached by no design, and transfer to Cas12a fails by five to six nucleotides in a way that localizes to the fusion junction rather than to reach, sterics or substrate. Tether geometry, therefore, bounds where a fused deaminase can act without predicting where it does so, making attachment site rather than linker length the effective design variable.

## INTRODUCTION

Much of the genome-editing toolkit works by tethering a programmable DNA-binding module that locates a target and exposes a nucleic acid substrate, with an effector domain attached to the module that acts on the substrate. In base editing, the effector is a cytidine or adenosine deaminase, and the substrate is the non-target strand displaced within the R-loop.^1,2^ The activity window, meaning the set of protospacer positions converted, determines whether an editor is therapeutically usable, because conversion of neighboring bases is the principal barrier to clinical application.^3,4^ Window engineering has proceeded continuously since the first base editors were reported. Lengthening the linker between rAPOBEC1 and Cas9 broadened the window,^1^ whereas short rigid linkers narrowed it.^5^ Inserting the deaminase into unstructured loops of the protospacer adjacent motif (PAM)-interacting and RuvC domains,^6^ replacing the HNH domain,^7^ and circularly permuting the scaffold^8^ each relocated it. Cas12a-based editors place the window at entirely different coordinates.^9^ Domain insertion in Nme2Cas9 was undertaken specifically to position the deaminase nearer the displaced strand,^10^ and systematic linker-length variation has recently been used to reduce bystander editing in plant and human cells.^11^

The reasoning behind these efforts was stated explicitly: the physical distance between the Cas domain and the deaminase, together with the rigidity of the connection, determines window width.^5^ That proposition has never been evaluated quantitatively for R-loop editors. Existing predictive models learn a mapping from guide sequence to editing outcome for a fixed editor.^12–17^ They interpolate across targets but cannot extrapolate to architectures absent from their training data, which is precisely the extrapolation an architecture-design question demands. The geometric argument has been made quantitative in mitochondrial editors built on transcription activator-like effector (TALE) proteins. Cryo-electron microscopy of DdCBE identified the structural determinants of its window and supported a model predicting editing outcomes across designed spacer lengths.^18^ Related work used de novo design to construct a rigid TALE– deaminase interface, confirming structurally that spatial architecture restricts the window.^19^ These systems differ from R-loop editors in two respects: the substrate is a rigid duplex and the engineered variable is the DNA spacer; here, the substrate is a flexible displaced strand and the variable is the protein tether. Effective concentration in tethered catalysis is a validated formalism for relating linker properties to reaction geometry,^20–22^ and it has not previously been applied to genome editors. We use it here to ask what a tethered deaminase would do if it sampled the substrate under linker statistics, steric exclusion and R-loop geometry alone. Agreement with the measurement would indicate that the geometry is sufficient, and disagreement would indicate what else is required.

## METHODS

### Model

#### Frames

The anchor frame is the coordinate of the fusion attachment residue on the substrate-bound structure, and it covers amino-terminal, carboxy-terminal, internal-insertion and domain-replacement anchors. The effector frame is the active-site position relative to the deaminase’s own fusion terminus, measured from deposited structures.

#### Linker Ensemble

Discrete worm-like chains with rise and persistence length assigned by composition class, grown from the anchor along departure directions drawn from a class-specific cone; 20,000 chains per draw, resampled to a fixed surviving count. Persistence values were XTEN 6.4 Å (range 4.5 to 6.4), glycine–serine 4.5 Å (range 4.5 to 4.8), polyproline 44.0 Å (range 20 to 130) and α-helical 180.0 Å (range 150 to 250). The XTEN value is bracketed by measurements at defined glycine content;^23^ the glycine–serine value is a worm-like-chain fit to Förster resonance energy transfer across GlyGlySer repeats;^24^ the polyproline value is a worm-like-chain fit to single-molecule FRET,^25^ and the band it carries runs from an apparent 20 to 40 Å under FRET^26^ to 90 to 130 Å for the all-trans form under all-atom simulation;^27^ and the α-helical value lies between light scattering on α-helical poly-L-lysine at 150 to 210 Å^28^ and an NMR measurement on a single α-helical domain at 224 Å.^29^ No direct measurement of XTEN persistence is available, and the assigned value lies at the stiff end of the bracket on the basis that its two proline residues restrict backbone flexibility. Rigid classes are assigned wide departure cones.

#### Substrate Ensemble

The displaced non-target strand is modeled as a discrete worm-like chain pinned at nucleotides held by base pairing, determined by direct distance measurements at a 12.0 Å threshold, and sampled as bridges between pins; 8000 substrate samples per entry.

#### Steric Exclusion

A union of spheres over the protein with uniform grid hashing. Conformations that intersect the excluded volume are discarded during chain growth.

#### Occupancy and Link

The linker field is convolved with the substrate ensemble within a capture radius and multiplied by the deaminase’s intrinsic motif preference, yielding a per-position capture value. The predicted profile follows from a saturating link,

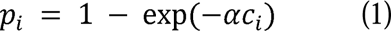

where *p_i_* is the predicted conversion propensity at protospacer position *i*, *c_i_* is the capture value at that position, and α is the link steepness. Mode is evaluated on the pre-link-capture profile, which is analytically equivalent because the link is strictly increasing and avoids loss of precision at high capture probability. Window spans are evaluated on linked values. Profiles whose peak ties across more than half their positions are excluded from reported rates.

#### Fitting

Three parameters over a 10 × 20 × 25 grid, on two entries, using the full per-position profile shape with the multiplicative constant solved by least squares. Peak normalization was not used: on synthetic profiles, it biases the recovered link steepness by approximately one-third and places the generating value outside the recovered interval.

#### Scoring

Predictions are scored with each position’s own anchor withheld, restricted to interior anchors, because removing the sole anchor on one side leaves the strand unbounded instead of interpolated. All reported values use a single fixed random seed, and sensitivity to seed is reported separately.

### Structures

SpCas9 geometry is taken from PDB 5F9R,^30^ which resolves displaced-strand positions 12 to 20, so that the window region is predicted rather than constrained. Cas12a geometry is taken from PDB 6I1K (FnCas12a),^31^ which resolves positions 1 to 13 and 19 to 21. PDB 6VPC (ABE8e)^32^ is used only as a steric consistency check and as an observation of an engaged complex, and it constrains no ensemble. PDB 5XUS (LbCas12a)^33^ is used only to evaluate the effect of the FnCas12a substitution. Residue numbering in the 6VPC coordinate file is offset by 197 from canonical SpCas9 numbering, because the deposited construct numbers the amino-terminal deaminase and its linker ahead of the Cas9 domain; all 6VPC residue indices were converted to canonical numbering before use. Protospacer numbering follows the convention used in each family’s literature: from the PAM-distal end for Cas9 and from the PAM-proximal end for Cas12a.^1,9^ No signed quantity is pooled across families without distance conversion.

### Statistics

Confidence intervals are estimated from the source studies, not the entries, because entries cluster by laboratory, study, deaminase, and scaffold. Intervals are studentized wild-cluster bootstrap with 1999 resamples, and 95% coverage.^34^ Sets containing a single cluster are reported as point estimates without intervals.

### Corpus Construction and Data Curation

Entries were read from primary figures, supplementary tables or deposited construct maps, with the source panel or table recorded. Entries carry labels distinguishing linker variation from deaminase variation and scaffold modification, and they record readout class: amplicon sequencing, in vitro deamination, Sanger deconvolution or functional selection. Entries not measured by amplicon sequencing are used for evaluation only and contribute neither to fitting nor to width analyses.

## RESULTS

### A Geometric Model of Tethered Deaminase Positioning

The editor is represented in the frame of the residue to which the deaminase is fused (Figure 1A). Three distributions are combined. The linker is sampled as a discrete worm-like chain with a persistence length assigned by composition class, grown from the anchor along a departure direction drawn from a class-specific cone, with the deaminase placed at the far end in a uniformly sampled orientation. Only the magnitude of the active-site offset enters the prediction, because a flexible linker leaves the domain rotationally decorrelated from the anchor. The displaced non-target strand is sampled as a second discrete worm-like chain, pinned at the nucleotides held by base pairing and sampled as bridges between them. Steric exclusion is represented as a union of spheres over the protein, and conformations that intersect it are discarded during chain growth. Convolution of the linker field with the substrate ensemble at a given capture radius yields a per-position capture probability, which is multiplied by the deaminase’s intrinsic motif preference and passed through a saturating link to yield the predicted profile (eq 1, Figure 1B). Insertions carry an additional closure term that weights each conformation by the contribution of the return linker at the separation it must span. Three parameters are fitted: capture radius, displaced-strand persistence and link steepness. Fitting uses the full per-position profile shape rather than the window position, because the window position is invariant under any monotone link, and a position-only objective would leave the link unidentified. Window position is the quantity on which all subsequent predictions are evaluated. The parameters were fitted to two entries from a single previously reported profile series and held fixed thereafter (Figure 1C). Predicted and measured profiles for all 26 profile entries are given in Figure S1.

**Figure 1.**
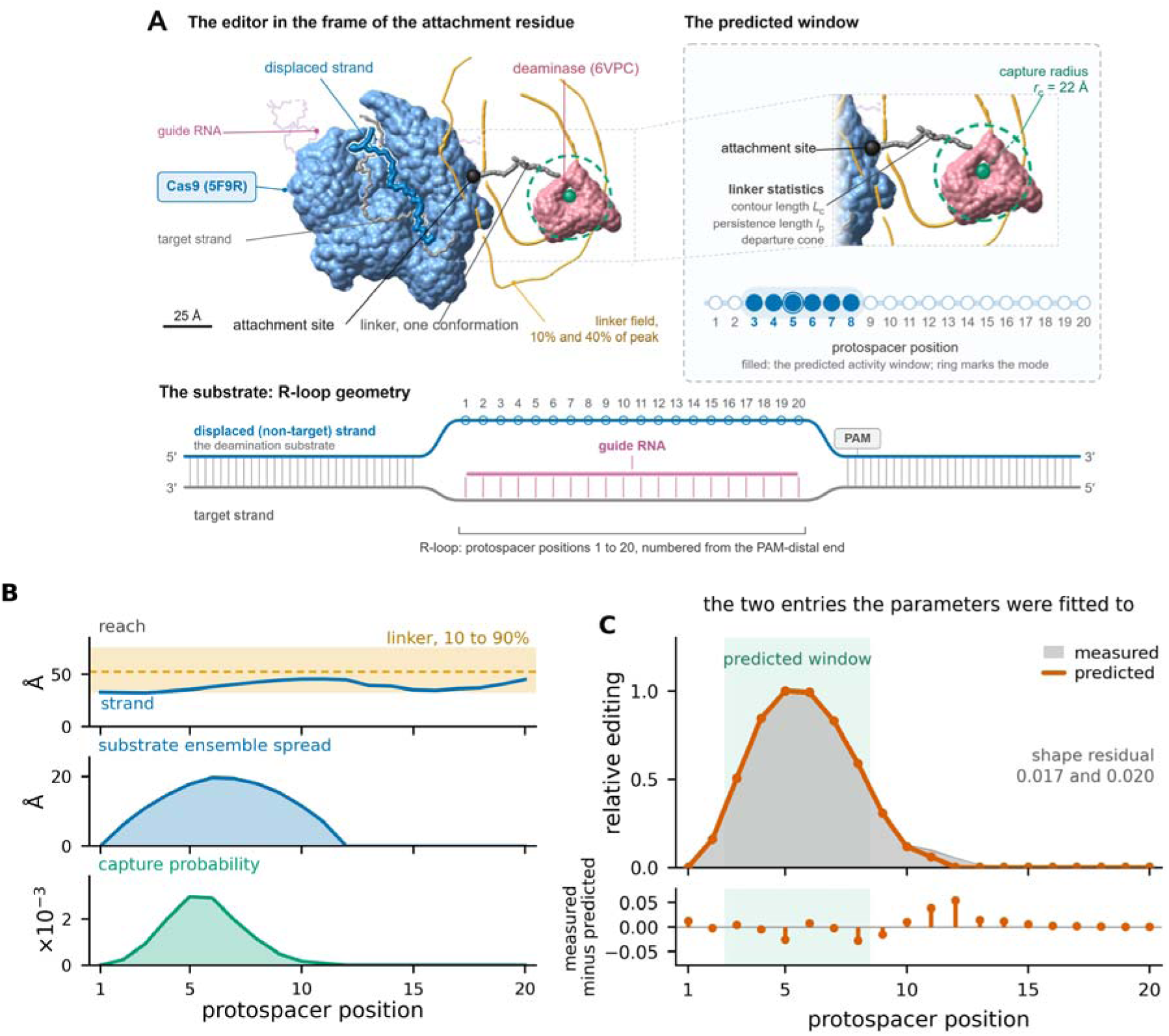
Geometric model of tethered deaminase positioning. Panels A and B are the sampled ensembles themselves, drawn at the frozen parameters, for the amino-terminal 32-residue XTEN entry to which the parameters were fitted. (A) The editor in the frame of the fusion attachment residue, the junction in close-up, and the R-loop substrate that the deaminase acts on. SpCas9, the guide, and both DNA strands are drawn from PDB 5F9R, and the deaminase from PDB 6VPC, as cited in Methods. The linker is one sampled conformation that clears the excluded volume and ends at the ensemble’s median reach; a flexible linker leaves the domain rotationally decorrelated, so its orientation is arbitrary and is taken as deposited. Orange contours enclose the sampled linker field at 10% and 40% of its peak density, and the dashed circle is the fitted capture radius about the active site. Filled circles mark the predicted activity window, protospacer positions 3 to 8, with a ring on the mode at position 5. Panel A is a prepared composite; every quantity in it is taken from the same run as panels B and C. (B) The three per-position quantities the convolution combines: the linker’s active-site reach, as its 10th to 90th percentile band and median, against the distance from the anchor to each strand position; the spread of the sampled strand about its own mean; and the capture probability the two give. (C) Predicted and measured per-position profiles for the two entries to which the parameters were fitted.

### Corpus

The corpus comprises 50 architectures drawn from seven studies (Table 1): 25 SpCas9, 18 SpRY, 4 SpG and 3 Cas12a; 43 with XTEN linkers, 4 with glycine-serine and 3 with polyproline; and 40 terminal fusions alongside 10 non-terminal anchors, the latter comprising PAM-interacting and RuvC insertions, HNH replacements and four circular permutants. Twenty-six entries carry a per-position profile and 15 a reported window. Linker lengths span 3 to 32 residues. Every entry is listed with its provenance in Table S1.

**Table 1.** Corpus composition by ortholog, linker class, anchor type, anchor site, readout class and measurement type, with linker length range and source counts.

| Category | Value | Count |
| --- | --- | --- |
| Ortholog | SpCas9 | 25 |
| Ortholog | SpRY | 18 |
| Ortholog | SpG | 4 |
| Ortholog | Cas12a | 3 |
| Linker class | XTEN | 43 |
| Linker class | Glycine–serine | 4 |
| Linker class | Polyproline | 3 |
| Anchor | Terminal | 40 |
| Anchor | Non-terminal | 10 |
| Anchor site | Amino terminal | 40 |
| Anchor site | PI domain, amino side | 3 |
| Anchor site | HNH domain, amino side | 2 |
| Anchor site | RuvC domain, amino side | 1 |
| Anchor site | CP1012 | 1 |
| Anchor site | CP1028 | 1 |
| Anchor site | CP1041 | 1 |
| Anchor site | CP1249 | 1 |
| Readout class | Amplicon sequencing | 38 |
| Readout class | Sanger deconvolution | 5 |
| Readout class | Functional selection | 4 |
| Readout class | In vitro deamination | 3 |
| Measurement | Per-position profile | 26 |
| Measurement | Reported window | 15 |
| Linker length | Range | 3 to 32 residues |
| Entries | Total | 50 |
| Studies | Contributing entries | 7 |
| Studies | Architecture sources examined | 16 |

Linker length is recorded in residues read from construct maps, not by linker name, because “XTEN” denotes both sixteen-residue and thirty-two-residue linkers in the literature.^1,17^ Of the sixteen architecture sources examined, six met the criteria for inclusion; the remainder are unavailable, lack tether variation, or report insertion residues or linker lengths only within figure panels (Table S2). Twenty-six entries derive from a single study, library and assay, and every architecture in that source peaks at position 5 or 6. Rates computed over the full corpus, therefore, reflect that concentration, so the analyses below use the architecture entries from the remaining studies wherever held-out performance is at issue.

### Parameter Identifiability

Refitting across five subsets that span an order of magnitude in size returns a capture radius between 21 and 25 Å, a displaced-strand persistence between 8 and 40 Å, and a link steepness between 90 and 1493 (Table S3). The capture radius is therefore identified, and the other two are not. Fit loss rises from 0.015 on two profiles to 0.975 on twenty-six, which indicates that a single parameter set does not describe all profiles in the corpus. The residual is structured by deaminase and not by delivery: after least-squares scaling, the root-mean-square shape residual is 0.039 across the thirteen ABE8e entries, on which the parameters were fitted, and 0.067 across the thirteen ABEmax entries (Figure S1). Both unidentified parameters govern profile shape. Across all five parameter sets, the median mode spread is 1 nucleotide, and the maximum is 2. The two Cas12a entries are set aside here because their profiles are degenerate across three of the five parameter sets, and their mode shifts by seven positions under the other two. Hit rates within one nucleotide range from 0.82 to 0.85 over the corpus and from 0.46 to 0.54 over the architecture entries. Predicted window positions are therefore insensitive to the parameters that the data do not constrain. Varying the XTEN persistence assignment across its literature-supported range of 4.5 to 6.4 Å^23,24^ changes the predicted mode by at most one nucleotide for all 38 of the 43 entries in that class that the walk scores. The hit rate moves by four percentage points across the band, entirely because the two Cas12a entries drop out of the denominator at its stiff end (Table S4, Section S1).

### Contour Length Does Not Move the Predicted Window

Length series provide the most direct test of a tethering model, and the expected direction is unambiguous: a longer tether should reach further from the anchor. Across an amino-terminal XTEN linker from 3 to 48 residues, evaluated at unfitted defaults, the predicted peak remains at position 5 or 6 for both ABE8e and APOBEC1, and the center of mass varies by 1.4 nucleotides without ordering by length (Figure 2A). Linker length is therefore not the variable that positions the window in this model.

**Figure 2.**
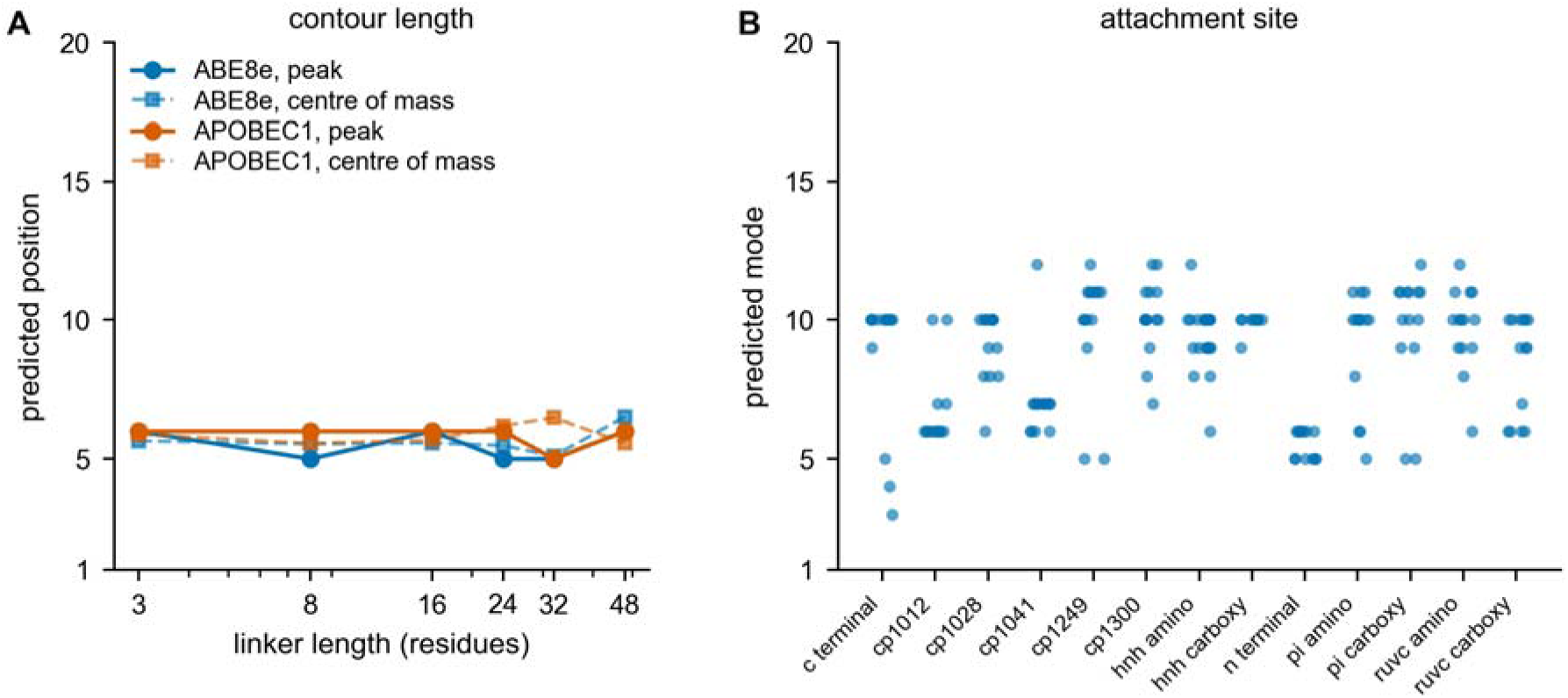
Contour length does not move the predicted window; the attachment site does. (A) Predicted peak position and center of mass against linker length from 3 to 48 residues for ABE8e and APOBEC1, evaluated at unfitted defaults. (B) Predicted mode of the 185 buildable candidate architectures, by attachment site. Both panels share a protospacer position axis from 1 to 20.

### Attachment Site Bounds the Accessible Set

Evaluating the model across 13 attachment sites, three composition classes, and seven contour lengths ranging from 4 to 48 residues yields 273 candidate architectures, of which 185 are sterically buildable. Their predicted modes fall on protospacer positions 3 to 12 (Figure 2B, Figure 3A). Across five independent sampling seeds, the union of accessible positions is 3 to 12, the intersection is 5 to 11, and positions 1, 2 and 13 to 20 are the predicted mode of no candidate at any seed (Figure 3B, Table S4). Boundary variation tracks candidate density: position 10 carries 63 to 80 candidates across seeds and position 6 carries 27 to 31, whereas position 3 is reached by a single candidate at two seeds and by none at the other three. A single-linker design on the intact scaffold, therefore, cannot place the peak outside positions 3 to 12; it places the peak within 5 to 11 across all sampling conditions. The candidate set comprises single-linker fusions on the unmodified scaffold in the XTEN, polyproline and helical classes, anchored at either terminus, at five circular-permutation termini and at six domain-boundary sites. Every candidate carries one outgoing connection and no return path, so the bound applies to designs that make a single connection to an intact scaffold rather than to insertions evaluated with their return linker. One corpus entry falls outside the bound without contradicting it: an HNH-replacement construct predicted at position 18 uses the glycine–serine class, carries a return linker, and sits on an HNH-excised scaffold whose excluded volume differs from that of the candidate set. The bound concerns the predicted mode, not the window span. All twelve tether-varying SpCas9-family entries with a reported window have their center within positions 3 to 12, across five studies, although two windows extend beyond the band as spans (Figure 3C). This is a consistency check rather than an independent test, since these entries are not held out from the accessible set. Because pinning inflates capture toward the PAM-proximal end, the model errs toward including positions rather than excluding them, and the inaccessible set is correspondingly more secure.

**Figure 3.**
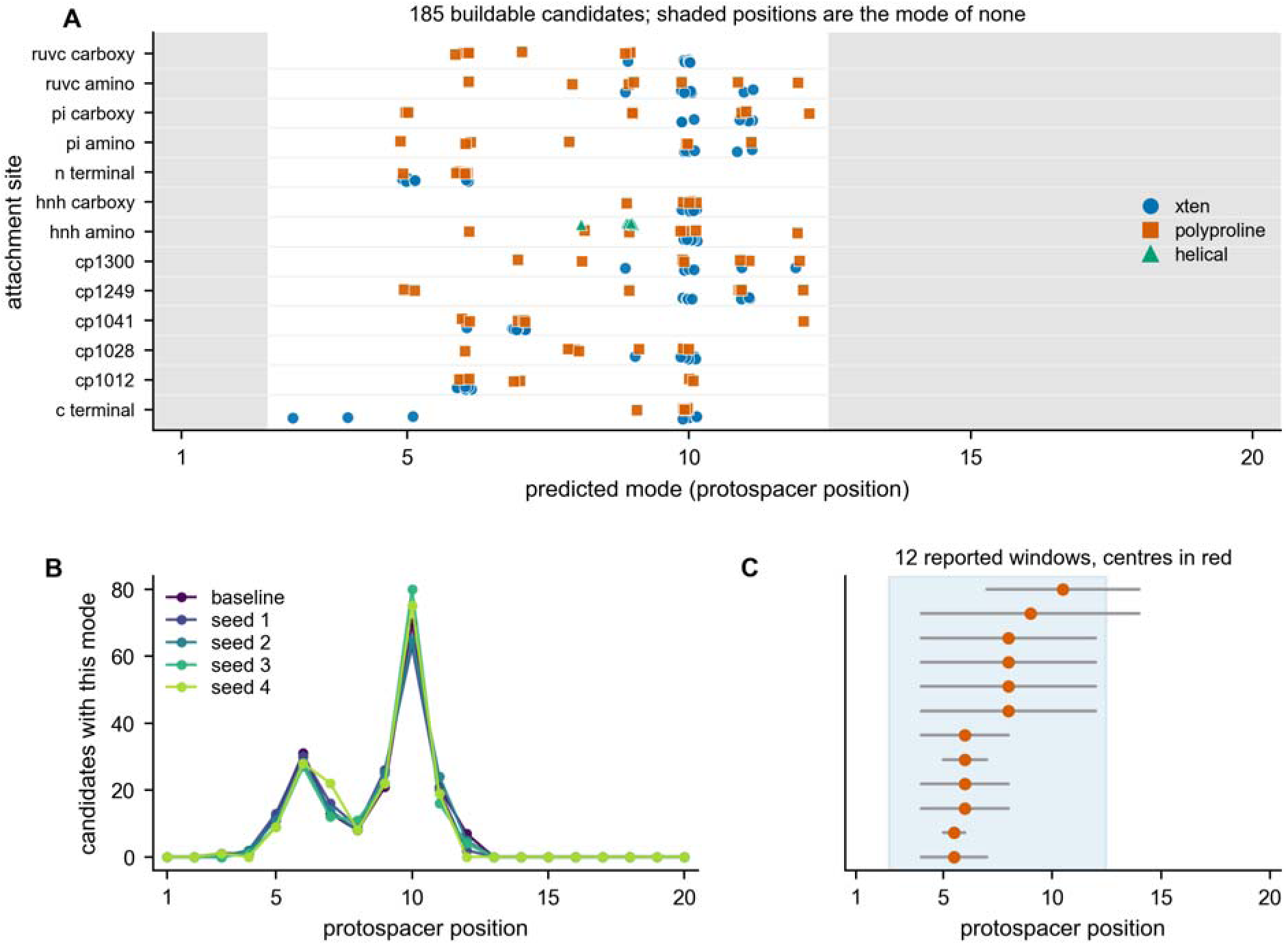
The accessible band. (A) Predicted mode of each buildable candidate across 13 attachment sites, three linker classes and seven contour lengths; shaded positions are the mode of no candidate. (B) Number of candidates with each predicted mode, at the configured seed and at four others. (C) Reported windows of the twelve tether-varying SpCas9-family entries that report one, with centers marked, against the accessible band. Panel C is a consistency check and not an independent test.

### Connector Requirements for Inaccessible Positions

For each inaccessible position, we computed the span, direction cone and steric admissibility that a connector would require, measured from the amino terminus (Table 2). Spans range from 40.8 Å for position 1 to 56.0 Å for position 20, corresponding to 16 to 23 polyproline residues or 28 to 38 residues of α-helix. Direction cones at 68% enclosure range from 26° to 44°, and 24% to 36% of departure directions from the junction are sterically allowed. The PAM-proximal positions impose a tighter constraint, and any connector that reaches them must be oriented and sized accordingly.

**Table 2.** Connector requirements for the positions no candidate design reaches: required span, direction cone, steric admissibility and residue equivalents.

| Position | Span (Å) | Span range (Å) | Cone at 68% (°) | Allowed fraction | Polyproline (res.) | α-Helix (res.) |
| --- | --- | --- | --- | --- | --- | --- |
| 1 | 40.8 | 23.2 to 57.5 | 43 | 0.33 | 16 | 28 |
| 2 | 40.8 | 23.5 to 57.5 | 44 | 0.34 | 16 | 28 |
| 13 | 50.1 | 32.4 to 67.1 | 32 | 0.36 | 20 | 34 |
| 14 | 49.2 | 31.9 to 66.5 | 34 | 0.31 | 20 | 33 |
| 15 | 46.3 | 29.2 to 62.3 | 37 | 0.27 | 19 | 31 |
| 16 | 45.4 | 28.6 to 61.4 | 37 | 0.24 | 18 | 31 |
| 17 | 47.9 | 30.7 to 64.1 | 34 | 0.25 | 19 | 32 |
| 18 | 49.2 | 31.2 to 66.4 | 32 | 0.25 | 20 | 33 |
| 19 | 52.3 | 34.1 to 70.1 | 29 | 0.29 | 21 | 35 |
| 20 | 56.0 | 37.6 to 72.8 | 26 | 0.33 | 23 | 38 |

Span, required distance from the amino terminus of PDB 5F9R to the target position, at a capture radius of 22.0 Å; cone, direction cone at 68% enclosure; allowed fraction, sterically permitted fraction of departure directions from the junction; res., residue equivalents of the required span.

### Anchor Transfer

Eight anchor relocations were predicted with the parameters held fixed (Figure 4A). Seven of the eight move in the reported direction. Over the six SpCas9-family relocations, the mean absolute error is 1.58 nucleotides, and the model overpredicts the magnitude of relocation (mean +3.17 against a reported +2.92); over all eight relocations, the corresponding means are +2.88 and +1.94. Reported shifts cluster narrowly, with four of the six at +2.50 nucleotides, so these results establish that the model reproduces the existence and approximate magnitude of an attachment-driven relocation rather than discriminating among anchors. Among the four circular permutants, the model does not identify the broadest window: CP1028 and CP1041 are predicted to be equal-widest at five positions, compared with CP1012 at four and CP1249 at three.

**Figure 4.**
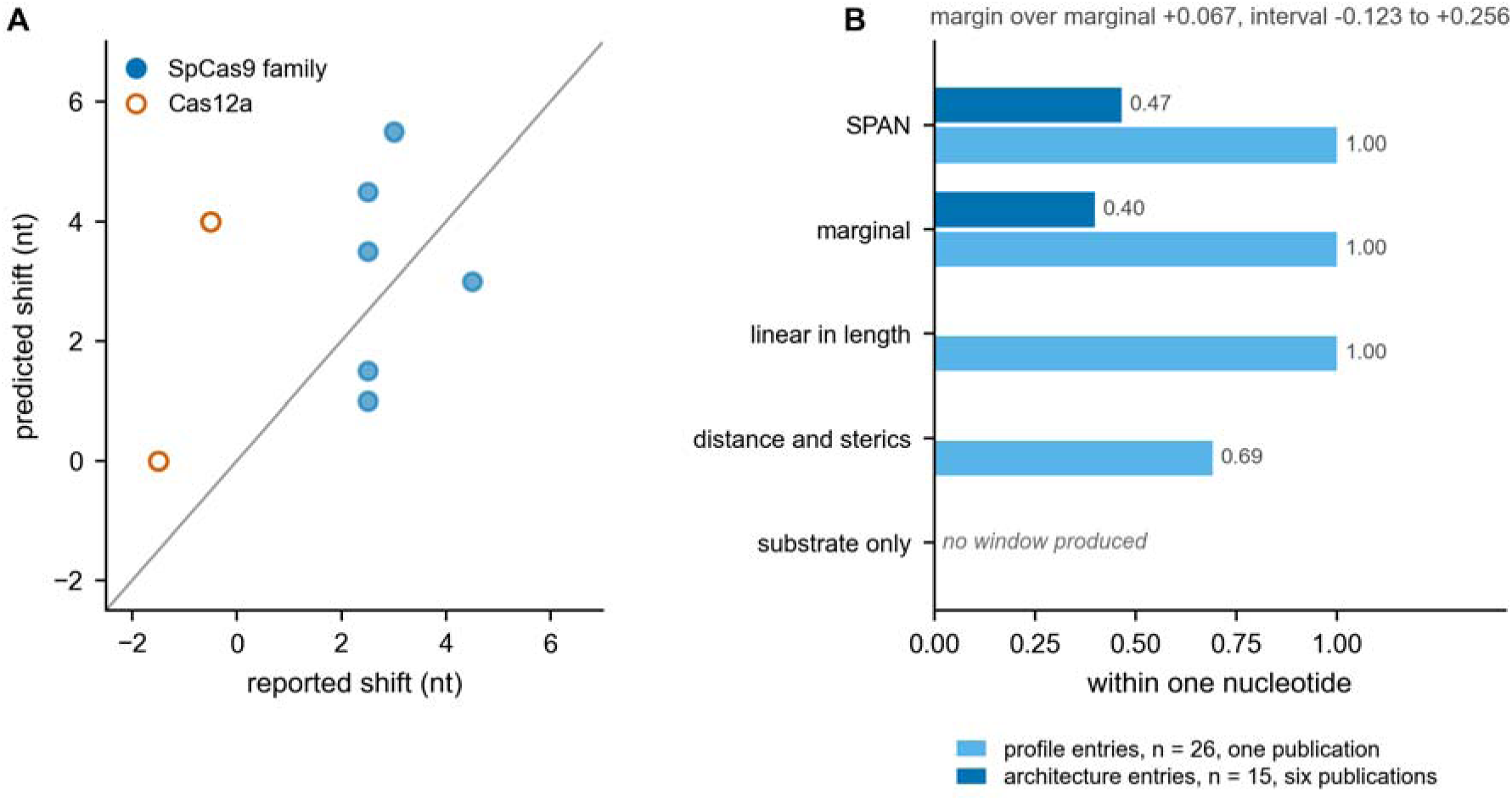
Anchor transfer and comparison against baseline predictors. (A) Predicted against the reported window shift for eight anchor relocations, with the identity line. (B) Hit rate within one nucleotide on the architecture entries, where a cluster bootstrap over six studies is possible, and on the profile entries, which come from a single study and therefore carry no interval.

### Predicted Window Position against Baselines

On the 15 held-out architecture entries, the model places the mode within one nucleotide in 0.467 of the entries, compared to 0.400 for a marginal baseline that predicts the fitted source mean at every entry. The margin is +0.067 (95% cluster bootstrap interval −0.123 to +0.256; Table 3, Figure 4B) and does not exclude zero. Under the same resampling scheme, six clusters do not reach 80% power at any simulated margin, and detecting a 25-point margin would require nine source studies (Section S2).

**Table 3.** Predicted window position against baseline predictors, on the architecture entries and the profile entries separately.

| Set | Predictor | n | Sources | Within 1 nt | Signed offsets | Margin (95% CI) |
| --- | --- | --- | --- | --- | --- | --- |
| Architecture | SPAN | 15 | 6 | 0.467 | −6 to +8 | Reference |
| Architecture | Marginal | 15 | 6 | 0.400 | −4 to +0 | +0.067 (−0.123 to +0.256) |
| Profile | SPAN | 26 | 1 | 1.000 | −1 to +0 | NA |
| Profile | Marginal | 26 | 1 | 1.000 | +0 to +1 | NA |
| Profile | Linear in length | 26 | 1 | 1.000 | +0 to +1 | NA |
| Profile | Distance and sterics | 26 | 1 | 0.692 | +1 to +2 | NA |
| Profile | Substrate only | 0 | 1 | NA | NA | NA |

Within 1 nt, predicted mode falling within one nucleotide of the measured window position; Sources, number of source studies contributing entries; CI, confidence interval. Intervals resample source studies, and sets containing a single cluster are reported without intervals. Reference, the predictor against which margins are computed; NA, quantity not defined; the substrate-only predictor produced no window.

On the 26 profile entries, the model, the marginal baseline and a linear-in-length predictor each place the mode within one nucleotide for every entry; a distance-plus-steric-exclusion predictor achieves 0.69, and a substrate-only model produces no window at all. The model does not improve on a constant predictor on any subset of this corpus. Separating the results by family, the SpCas9 family is within one nucleotide for 0.58 of 12 entries, with offsets between −2 and +8, and Cas12a for none of the three, with offsets of −5 and −6. The largest positive offset is observed for the HNH-replacement construct, whose predicted mode lies outside the single-linker-accessible set. Window width is not reproduced, and the errors run in both directions: +0.81 nucleotides on the profile source and −3.53 nucleotides on the architecture entries, spanning −10 to +3 (Table S5). Link steepness, the parameter that governs width, is the least constrained of the three.

### Transfer to Cas12a Fails at the Attachment Site

The three Cas12a entries come from a single study,^35^ and the construct was read from its deposited plasmid (Addgene 193640): human APOBEC3A, then the sixteen-residue linker SGSETPGTSESATPES, then LbCas12a, with the deaminase amino-terminal. At that anchor, the model places the ABE8e mode at position 4, against a reported window of 8 to 12. Scored from the carboxy terminus, every contour length returns a mode of 9 or 10 and a window of 9 to 12, against reported windows of 6 to 15, 6 to 12 and 8 to 12. Since the substrate ensemble, the steric map and the fitted parameters are common to both scorings, none of these components accounts for the discrepancy. Four further explanations were examined and excluded: reach, capture radius, structural substitution and seed sensitivity (Figure 5). Sampling the sixteen-residue linker with its deaminase gives a mean reach of 42.4 Å for dCas12a-ABE8e and 38.6 Å for dCas12a-A3A, against the 38 to 40 Å range reported, with 45% to 55% of the ensemble weight beyond 40 Å. The contour length is 57.6 Å, and the active site lies approximately 21 Å beyond the chain terminus. The model assigns non-zero probabilities to positions 8, 9, and 10, so the discrepancy lies in the weighting, which peaks at position 4, with 9 times the probability of position 8.

**Figure 5.**
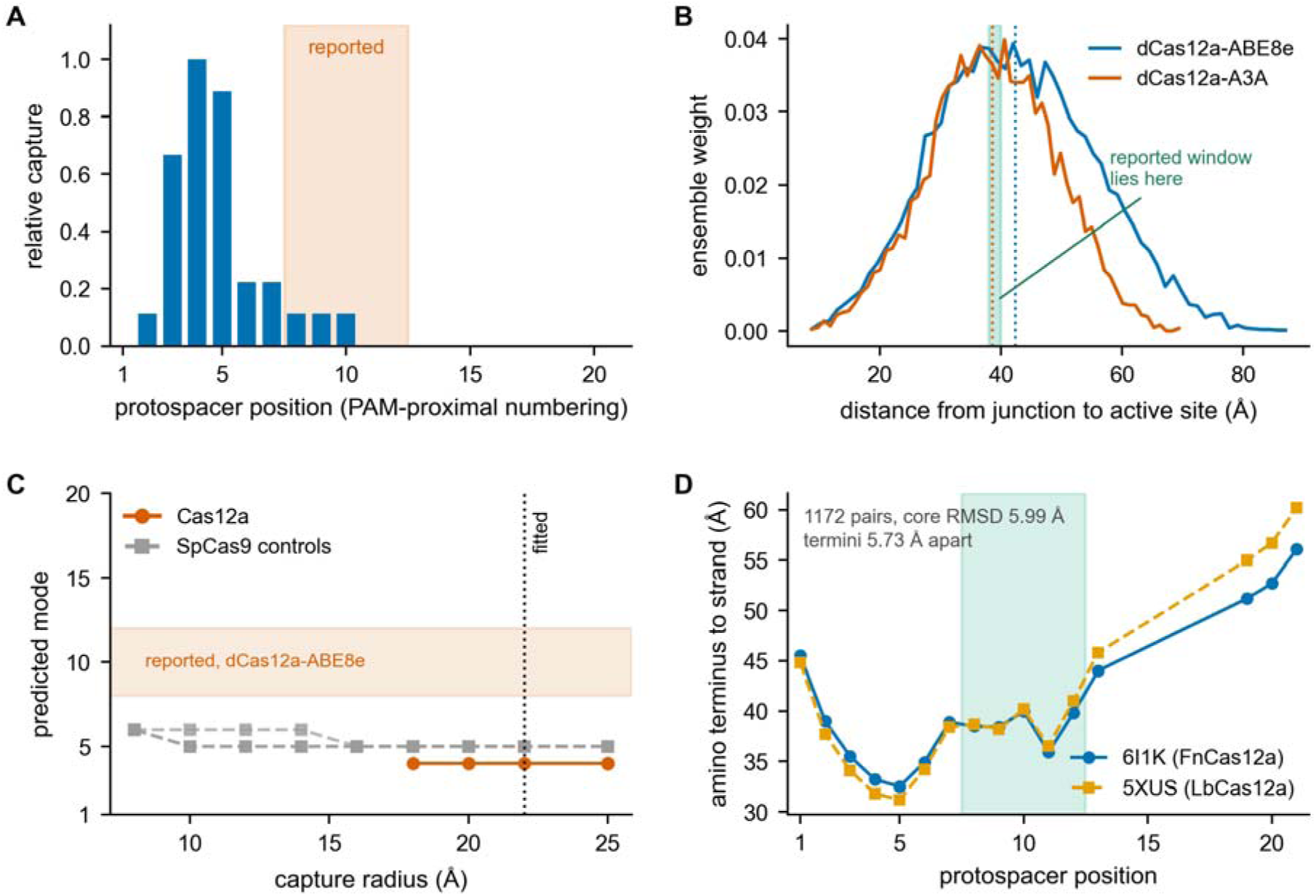
The Cas12a discrepancy localizes to the junction. (A) Predicted capture profile at the verified amino-terminal anchor, with the reported window shaded. (B) Distribution of distances from the fusion junction to the sampled active site for the sixteen-residue linker, with the distance at which the reported window lies marked; dotted lines mark the means. (C) Predicted mode against capture radius from 8 to 25 Å for the three Cas12a entries and three SpCas9 controls, with all other parameters held fixed. (D) Distance from the amino terminus to each displaced-strand position in FnCas12a and in LbCas12a after superposition, with the reported window shaded.

Sweeping the capture radius from 8 to 25 Å with all other parameters held fixed, no Cas12a entry has its mode fall within its reported window at any radius: no entry produces a window below 18 Å, and wherever one is produced at or above 18 Å, the mode is 4 and does not vary, while the two APOBEC3A entries produce none until 25 Å. Three SpCas9 controls remain within one nucleotide across all nine radii. When the FnCas12a structure used for geometry is superposed onto an LbCas12a structure at the same PAM (1172 residue pairs, 5.99 Å core root-mean-square deviation), the amino termini lie 5.73 Å apart, and their distances to the substrate over the reported window agree to within 0.5 Å. The Cas12a offset is −5.667 nucleotides at every seed tested, although its point value varies between positions 4 and 7, so the magnitude is reported as a range and not as a single value. A sixteen-residue amino-terminal tether therefore reaches the reported positions but is predicted to act predominantly elsewhere. Every component of the model except the anchor has been excluded, so the discrepancy localizes to what happens at the junction rather than to reach, sterics or the substrate.

### The Displaced Strand Is Not Recoverable from Tether Statistics

Sampling the displaced strand against the protein without coordinate information, the median distance between the sampled and deposited paths is 24.70 Å in the SpCas9 structure and 9.58 Å in the Cas12a structure at residue resolution. Refining the steric representation from residue centers to individual atoms, from 1508 to 13,256 spheres in the SpCas9 structure and from 1359 to 12,311 in the Cas12a structure, gives 22.78 Å and 10.62 Å, respectively (Figure 6).

**Figure 6.**
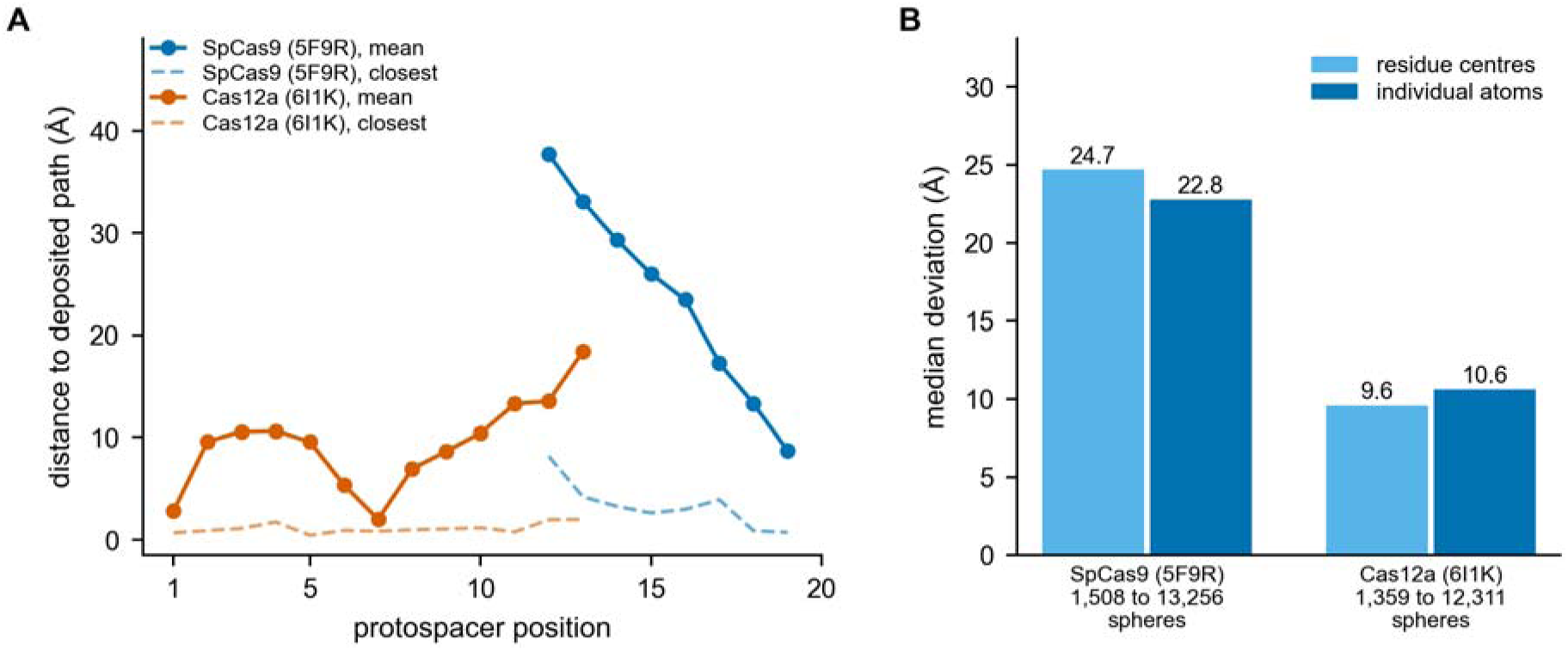
Displaced-strand position is not recoverable from tether statistics. (A) Mean and closest distance between the sampled strand ensemble and the deposited path, by protospacer position, for both structures. (B) Median deviation at residue and at atomic steric resolution.

Strand position is therefore not recoverable from linker statistics and steric exclusion at the resolutions tested. Deposited coordinates are consequently entered into the model as input, and predictions are scored with each position’s own anchor withheld, since an ensemble constrained to a nucleotide’s position will reproduce it. Anchoring follows base pairing determined by direct measurement, not the set of nucleotides a given deposition resolves, so that the substrate ensemble contributes equivalently across orthologs.

### Robustness

Varying only the sampling seed, 12 of the 15 architecture entries return identical modes, and two vary by one nucleotide. The exception is Cas12a, where profile contrast, rather than conformation count, is responsible: stable entries peak between 0.39 and 1.00 after the link, and Cas12a entries peak between 0.06 and 0.18, so low-contrast profiles allow small fluctuations to move the argmax. Predicted modes for low-contrast profiles are therefore not quoted to single-nucleotide precision. The profile source varies in delivery across nine contexts. Architecture changes profile shape 2.58-fold more than the delivery context does. Center positions shift by 0.127 and 0.133 nucleotides, respectively, both an order of magnitude below the resolution at which predictions are evaluated. For synthetic profiles generated at known parameter values, the fitting procedure recovers the capture radius and persistence exactly and links steepness to within 6%. A wild cluster bootstrap holds its nominal 5% rejection rate at zero margin across cluster counts, whereas a percentile cluster bootstrap rejects 6% to 13% over the same range and 13% at the six clusters available here; the wild bootstrap is therefore used throughout. Pinned nucleotides do not move, with a maximum positional spread of 0.00 Å, and no ensemble weight lies within the excluded volume.

## DISCUSSION

Tether geometry bounds where a fused deaminase can act on an R-loop without predicting where it does. Across 185 buildable single-linker designs spanning thirteen attachment sites, three composition classes and a sixteen-fold range of contour length, the predicted peak never falls outside protospacer positions 3 to 12, and it lies within 5 to 11 across all sampling conditions.

Independent of this, contour length does not move the predicted window at all. Taken together, these findings indicate that linker length is not the design variable it has been treated as, and that attachment site is. The conclusion is consistent with the published studies: architectures that relocated windows substantially did so by inserting the deaminase into scaffold loops, replacing a domain, or permuting the protein, not by lengthening a linker.^6–8,10^ It also accounts for the narrow range of editing windows produced by a decade of linker engineering. The design implications are direct. Positions 1, 2 and 13 to 20 are inaccessible to any single-linker design on an intact scaffold, so reaching them requires an attachment site outside the thirteen swept here, a modified scaffold, or a rigid connector. We specify the span, direction and rigidity that such a connector would require. The direction constraint is as restrictive as the span: only a quarter to a third of departure directions from the junction are sterically permitted, so a connector must be oriented rather than merely extended. Table 2 gives the requirement in buildable units: a span of 41 to 56 Å, which is 16 to 23 polyproline residues or 28 to 38 α-helix residues, leaving the junction within a cone of 26° to 44°. A de novo helical bundle or a polyproline scaffold is the natural route, since both fix the departure direction that a flexible linker would otherwise randomize. Only seven of the 185 buildable candidates were helical; the rest of that class intersected the scaffold, so a rigid connector has to be designed around the excluded volume rather than substituted for a flexible one. Rigid-interface design has achieved comparable window restriction experimentally in TALE systems,^19^ and the present analysis derives the equivalent requirement from the substrate geometry of R-loop editors.

Window position agrees with measurement to approximately one nucleotide but does not improve on a constant predictor. That predictor is hard to beat because the accessible band is narrow and its center is crowded: the baseline predicts position 6 at every entry, and 32 of the 41 scored entries were measured at position 5 or 6. Six source studies cannot resolve a margin of the observed size, so the comparison does not show that the two predictors perform equally (Section S2). The limitation is also evident in the fit: loss rises 65-fold as the fitting set widens, and the two parameters that govern profile shape are unidentified over an order of magnitude. The model reproduces where a window is centered but not its width. Transfer to Cas12a fails in a way that isolates the junction. A sixteen-residue amino-terminal tether demonstrably reaches the reported positions, with roughly half its ensemble weight beyond 40 Å, and the model assigns non-zero probability there, yet it concentrates nine times more probability at position 4 than at position 8. Reach, capture radius, steric map, substrate treatment, fitted parameters, and the structural substitution used for geometry were each excluded in turn. The remaining variable is the junction: the manner in which a deaminase departs from an amino-terminal fusion in this scaffold appears to orient or extend it in ways that isotropic sampling does not capture. Two mechanisms would leave that signature, and the present data do not distinguish between them. The junction may impose an orientational preference on the departing domain, or the deaminase may make a specific contact with the Cas12a surface that a model that only excludes volume cannot represent, as it cannot attract. Direct evidence for the second exists in a different editor, where dimerization of TadA-8e and its juxtaposition to the Cas9–DNA complex govern deamination efficiency.^36^ An orientational term at the junction is the natural extension of the model. The finding that the position of the displaced strand is not recoverable from linker statistics and steric exclusion, a median of 22.8 Å in SpCas9, even at atomic resolution, has methodological consequences beyond this model. Structural coordinates must be treated as an input to a tethering model, not as something it can derive, and any prediction evaluated against them must withhold the coordinate being evaluated, since otherwise the ensemble reproduces what it was given. Tether variation is distributed across a small number of studies spanning mammalian cell culture and gene therapy, yeast selection and purified-protein biochemistry, using four distinct readout classes. We deposit 50 entries from seven studies, with labels distinguishing tether variation from deaminase and scaffold variation, and the recorded readout class.

Half the corpus comes from a single study, and the number of independent architectural sources is smaller than the analysis ideally requires, which is why the baseline comparison is underpowered. Rigid linkers carry a wide departure cone, because the direction in which a rigid connector leaves a junction is determined by backbone geometry that available structures do not resolve; rigid-class predictions therefore carry correspondingly wide uncertainty. The insertion-closure weights conformations by the contribution of the return linker without evaluating the return path against the protein, so it bounds reach rather than treating it fully. The accessibility bound applies to single-linker designs on intact scaffolds and does not extend to insertions evaluated with their return linker, to domain replacements, or to scaffolds from which a domain has been removed. Window width is not reproduced, indicating that the model omits physics to which the predicted mode is insensitive; this limits confidence in extending it to deaminases or scaffolds that differ from those examined here. The substrate ensemble is a worm-like chain under steric exclusion alone. It carries no sequence-dependent base stacking, no attraction of any kind to the protein the strand runs along, and no breathing at the pinned nucleotides. Each would bias the path, and their absence is a candidate explanation for the residual 22.8 Å; a substrate model that both attracts and excludes is the obvious test.

## CONCLUSIONS

Tether geometry bounds where a fused deaminase can act on an R-loop, but does not predict where it does. Across 185 buildable single-linker designs spanning thirteen attachment sites, three composition classes and a sixteen-fold range of contour length, the predicted peak never falls outside protospacer positions 3 to 12 and lies within 5 to 11 across all sampling conditions, whereas contour length does not move the predicted window at all. Attachment site, not linker length, is therefore the variable that positions the window, a conclusion consistent with the published studies, in which the architectures that relocated windows did so substantially by domain insertion, domain replacement, or circular permutation. Positions 1, 2, and 13 to 20 are inaccessible to any single-linker design on an intact scaffold. Reaching them requires a span of 41 to 56 Å, equivalent to 16 to 23 polyproline residues or 28 to 38 α-helix residues, held within a departure cone of 26° to 44°, so a connector must be oriented as well as extended. Predicted window position agrees with measurement to approximately one nucleotide, but does not improve on a constant predictor on a corpus of this size, and window width is not reproduced. The failure of transfer to Cas12a localizes to the fusion junction rather than to steric effects, the substrate, or the fitted parameters, thereby identifying an orientational term at the junction as the extension the model now requires.

## DATA AND SOFTWARE AVAILABILITY

### Data

All data required to reproduce the results reported here are openly available. The corpus of 50 architecture entries, the source register with the reason for each exclusion, the fitted parameter sets with their losses, the seed and persistence sensitivity results, and the per-entry predicted and measured window positions and widths are provided as Supporting Information (Tables S1 to S5) and in machine-readable form in the code release, which is openly available at https://github.com/ahmedanees-m/span-editor and archived at Zenodo under DOI 10.5281/zenodo.22736738.^37^ Structural coordinates are obtained from the Protein Data Bank under accessions 5F9R, 6I1K, 6VPC and 5XUS. The verified LbCas12a base-editor construct was read from Addgene plasmid 193640.^38^ Sampled conformational ensembles are not deposited because they regenerate deterministically from the recorded random seed using the provided code. No new experimental data were generated in this study.

### Software Developed in This Work

SPAN v0.1.0, the sampling and scoring code that produced every result reported here, is released under the MIT license at https://github.com/ahmedanees-m/span-editor and is archived at Zenodo under DOI 10.5281/zenodo.22736738.^37^ It contains the corpus and its data dictionary, the parameter grids, configuration files, and the fixed random seeds; installation and reproduction instructions are in the repository README. The model is fully specified in the Methods section, and the complete set of associated parameters is provided in machine-readable form in the repository.

### Third-Party Software

The analysis was run on Python 3.11.15 with NumPy 2.1.3, SciPy 1.14.1, and Biotite 1.0.1, all open-source and installed from the Python Package Index. The complete environment, with every version pinned, is specified in docker/requirements.txt and the Dockerfile in the code release; it is built and run under Docker 29.1.3. No commercial or licensed software was used.

## ASSOCIATED CONTENT

### Supporting Information

The Supporting Information is available free of charge at the ACS Publications website. Predicted and measured per-position profiles for all 26 profile entries; full corpus of 50 architecture entries with ortholog, anchor site, linker class and residue count, deaminase, readout class, reported window, confound label and the panel or table from which each value was read; source register of the studies examined, with role, status and the reason for each exclusion; fitted parameters and loss across the five fitting subsets, with the corpus hit rate under each parameter set; seed sensitivity, band edges and per-position candidate density at five sampling seeds, and mode and hit rate across the XTEN persistence band; per-entry predicted and measured window position and width with the signed offset; persistence length assignment for each linker composition class and parameter recovery on synthetic profiles; power analysis for the baseline comparison and calibration of the wild cluster bootstrap against the percentile cluster bootstrap (PDF) The same five tables as sheets of one workbook, in machine-readable form, with a dictionary defining every column and the complete twenty-one-column corpus record that the printed Table S1 abridges (XLS)

## AUTHOR INFORMATION

### Authors

Anees Ahmed Mahaboob Ali, School of Bio Sciences and Technology, Vellore Institute of Technology, Vellore, Tamil Nadu, India;

## Author Contributions

A.A.M.A. and E.J.R.N. conceived and designed the study. A.A.M.A. developed the model, wrote the code, assembled the corpus and performed all analyses. E.J.R.N. supervised the work. Both authors interpreted the results, wrote the manuscript and approved the submitted version.

## Notes

The authors declare no competing financial interest, and no commercial or financial relationships exist that could be construed as a potential conflict of interest.

## ACKNOWLEDGMENTS

This research did not receive any specific grant from funding agencies in the public, commercial or not-for-profit sectors. The authors gratefully acknowledge Vellore Institute of Technology, Vellore, India, for access to computational resources and for the institutional support under which this study was carried out.

Generative AI was used during the preparation of this manuscript. Claude (Anthropic; models Claude Opus 5 and Claude Sonnet 5) was used for language editing of author-written text and for coding assistance in implementing and analyzing the SPAN code. No text, figures, or data were generated de novo by the tool. All AI-assisted output was reviewed, verified and edited by the authors, who take full responsibility for the content, accuracy and integrity of the manuscript.

## ABBREVIATIONS

ABE: adenine base editor
CP: circular permutant
DdCBE: DddA-derived cytosine base editor
FRET: Förster resonance energy transfer
PAM: protospacer adjacent motif
PDB: Protein Data Bank
PI: PAM-interacting
TALE: transcription activator-like effector
XTEN: extended recombinant polypeptide

