## Supplementary material for "Attachment site, not linker length, bounds tethered base editor windows": SI

This Supporting Information contains two sections of extended methods, one figure and five tables. Tables S1 to S5 are also supplied as sheets of the workbook SI\_data.xls, which opens with a dictionary defining every column.

### Section S1. Persistence length assignment and method validation

#### Assigned persistence lengths

Persistence length is an assigned input rather than a fitted parameter. The XTEN class carries 43 of the 50 corpus entries, including both entries the parameters were fitted to, and is therefore the largest single assumption in the model.

| Class | Assigned (Å) | Range carried (Å) | Basis |
| --- | --- | --- | --- |
| XTEN | 6.4 | 4.5 to 6.4 | No direct measurement of XTEN persistence is available. Bracketed by worm-like-chain fits to FRET across a glycine series, which give 4.5 Å at 33 percent glycine, 4.8 Å at 17 percent and 6.2 Å with none [van Rosmalen, Krom and Merks, Biochemistry 2017, doi 10.1021/acs.biochem.7b00902] |
| Glycine-serine | 4.5 | 4.5 to 4.8 | Worm-like-chain fit to FRET between fluorescent domains joined by GlyGlySer repeats [Evers, van Dongen, Faesen, Meijer and Merks, Biochemistry 2006, doi 10.1021/bi061288t], reproduced at 4.5 Å by van Rosmalen and colleagues |
| Polyproline | 44.0 | 20 to 130 | Reported estimates span an order of magnitude by method. A worm-like-chain fit to single-molecule FRET gives $44 \pm 9$ Å [Schuler, Lipman, Steinbach, Kumke and Eaton, Proc Natl Acad Sci U S A 2005, doi 10.1073/pnas.0408164102], which is the assigned value. An apparent persistence length of 20 to 40 Å has been estimated from FRET analyzed under a worm-like chain [Doose, Neuweiler, Barsch and Sauer, Proc Natl Acad Sci U S A 2007, doi 10.1073/pnas.0705605104], and all-atom simulation puts the all-trans form at 90 to 130 Å [Best, Merchant, Gopich, Schuler, Bax and Eaton, Proc Natl Acad Sci U S A 2007, doi 10.1073/pnas.0709567104]. Those two set the band. The early theoretical figure of 220 Å is not supported by the FRET work |
| Alpha-helical | 180.0 | 150 to 250 | Light scattering on α-helical poly-L-lysine gives 150 to 210 Å [Wilcox, Dingle, Saha, Hore and Morozova, Soft Matter 2022, doi 10.1039/d2sm00921h], and NMR on the single α-helical domain of myosin-VI gives $224 \pm 10$ Å [Barnes, Shen, Ying, Takagi, Torchia, Sellers and Bax, J Am Chem Soc 2019, doi 10.1021/jacs.9b03116]. The band is widened to 250 Å so that it brackets both |

The sixteen-residue XTEN linker, SGSETPGTSESATPES, is 19 percent glycine, which places it near the 4.8 Å measurement, but it also carries two proline residues in sixteen, and proline restricts backbone flexibility. The assigned value sits at the stiff end of the bracket on that basis, and the whole bracket is carried as a band.

#### Cost of the XTEN band

Walking the band at 4.5, 5.13, 5.77, 6.4 Å with the fitted parameters held and only the assigned input changed:

| Quantity | Value |
| --- | --- |
| Entries in the class that return a mode | 38 of 43 |
| Median mode change across the band | 1 nucleotide |
| Maximum mode change | 1 nucleotide |
| Entries whose mode does not move | 9 |

No predicted mode moves by more than one nucleotide anywhere in the band. Five of the 43 entries in the class are not scored by the walk and are therefore not counted: hnhx-ABEmax7.10-dHNH, bepigs-pi-3, bepigs-pi-8, bepigs-ruvc-16 and tan2019-xten-16. Of the 38 that are scored, two, the Cas12a A3A entries, return no mode at the stiff end of the band, which is why the hit rate quoted in the main text moves across it.

#### Parameter recovery on generated profiles

The fitting procedure was exercised on profiles generated by the forward model at known parameter values, with 16 synthetic architectures and Gaussian noise of standard deviation 0.08.

| Parameter | Generating value | Recovered |
| --- | --- | --- |
| Capture radius (Å) | 9.0 | 9.0 |
| Displaced-strand persistence (Å) | 14.0 | 14.0 |
| Link steepness | 75.80 | 79.74 |
| Saturation index | 3.000 | 3.156 |

Capture radius and persistence are recovered exactly. Link steepness is recovered to within 5.2 percent. The predicted mode matches the generating mode exactly for 16 of 16 synthetic architectures.

#### Profile scale convention

Profiles are compared as shapes with the multiplicative constant solved by least squares rather than by dividing by the peak. Peak normalization was tested on the same generated profiles and biases the recovered link steepness toward the linear end, placing the generating value outside the recovered interval. The least-squares convention is used throughout.

#### Source

`configs/assigned_inputs.yaml` records each assignment with its citation.
`analysis/synthetic_recovery.py` and `analysis/composition_sensitivity.py`; outputs in
`results/synthetic_recovery.json` and `results/composition_sensitivity.json`.

### Section S2. Power and bootstrap calibration for the baseline comparison

#### Purpose

The comparison between the model and the marginal baseline is made on the architecture entries, which come from six source studies. Because entries cluster by study, intervals resample studies rather than entries. This section gives the margin that design can resolve and the calibration of the two cluster bootstrap constructions.

#### Simulation design

Clusters carry 5 entries each, at a base hit rate of 0.45 and an intraclass correlation of 0.3. The nominal rejection rate is 0.05 at the 0.95 level. Each cell is 600 trials of 1500 bootstrap resamples, seed 20260101.

Two pairings are simulated. Under comonotonic pairing the model and the baseline succeed and fail together within a cluster, which is the favorable case and the one the real comparison resembles, since both predictors see the same entries. Under independent pairing their successes are unrelated, which is the conservative case.

#### Calibration at a zero margin

Rejection rate when the true margin is zero. A calibrated method rejects at the nominal 0.05.

| Method | Pairing | Studies | Rejection at zero margin |
| --- | --- | --- | --- |
| percentile | comonotonic | 6 | 0.000 |
| percentile | comonotonic | 9 | 0.000 |
| percentile | comonotonic | 12 | 0.000 |
| percentile | comonotonic | 16 | 0.000 |
| percentile | comonotonic | 30 | 0.000 |
| percentile | comonotonic | 60 | 0.000 |
| percentile | comonotonic | 100 | 0.000 |
| percentile | independent | 6 | 0.128 |
| percentile | independent | 9 | 0.098 |
| percentile | independent | 12 | 0.062 |
| percentile | independent | 16 | 0.070 |
| percentile | independent | 30 | 0.065 |
| percentile | independent | 60 | 0.053 |
| percentile | independent | 100 | 0.055 |
| wild | comonotonic | 6 | 0.000 |
| wild | comonotonic | 9 | 0.000 |
| wild | comonotonic | 12 | 0.000 |
| wild | comonotonic | 16 | 0.000 |
| wild | comonotonic | 30 | 0.000 |
| wild | comonotonic | 60 | 0.000 |
| wild | comonotonic | 100 | 0.000 |
| wild | independent | 6 | 0.062 |
| wild | independent | 9 | 0.047 |
| wild | independent | 12 | 0.047 |
| wild | independent | 16 | 0.058 |
| wild | independent | 30 | 0.052 |
| wild | independent | 60 | 0.048 |
| wild | independent | 100 | 0.055 |

The wild cluster bootstrap stays close to the nominal rate throughout, from 0.000 under comonotonic pairing to 0.062 at six clusters under independent pairing. The percentile construction reaches 0.128 under independent pairing, roughly twice its nominal rate at the smaller cluster counts. The studentized wild cluster bootstrap is therefore used for every interval reported.

#### Detectable margin

Smallest margin reaching 80 percent power, by method, pairing and number of source studies.

| Method | Pairing | Studies | Smallest detectable margin |
| --- | --- | --- | --- |
| percentile | comonotonic | 6 | 0.232 |
| percentile | comonotonic | 9 | 0.177 |
| percentile | comonotonic | 12 | 0.133 |
| percentile | comonotonic | 16 | 0.101 |
| percentile | independent | 6 | 0.378 |
| percentile | independent | 9 | 0.320 |
| percentile | independent | 12 | 0.284 |
| percentile | independent | 16 | 0.247 |
| wild | comonotonic | 6 | not reached in the simulated range |
| wild | comonotonic | 9 | 0.251 |
| wild | comonotonic | 12 | 0.165 |
| wild | comonotonic | 16 | 0.113 |
| wild | independent | 6 | not reached in the simulated range |
| wild | independent | 9 | 0.375 |
| wild | independent | 12 | 0.321 |
| wild | independent | 16 | 0.263 |

Under the reporting method and the favorable pairing, six studies do not reach 80 percent power at any margin simulated: at the largest margin tested, 0.50, power is 0.79. At a margin of 0.075, close to the 0.067 actually observed, power is 0.055, which is the false-positive rate. Nine studies would be needed to resolve a margin of 0.25 and sixteen to resolve one of 0.11.

The null reported in the main text is therefore a property of the corpus size rather than evidence that the model and the baseline perform equally.

#### Source

analysis/power\_sim.py and analysis/bootstrap\_calibration.py; outputs in results/power\_\*.json and results/bootstrap\_calibration.json.

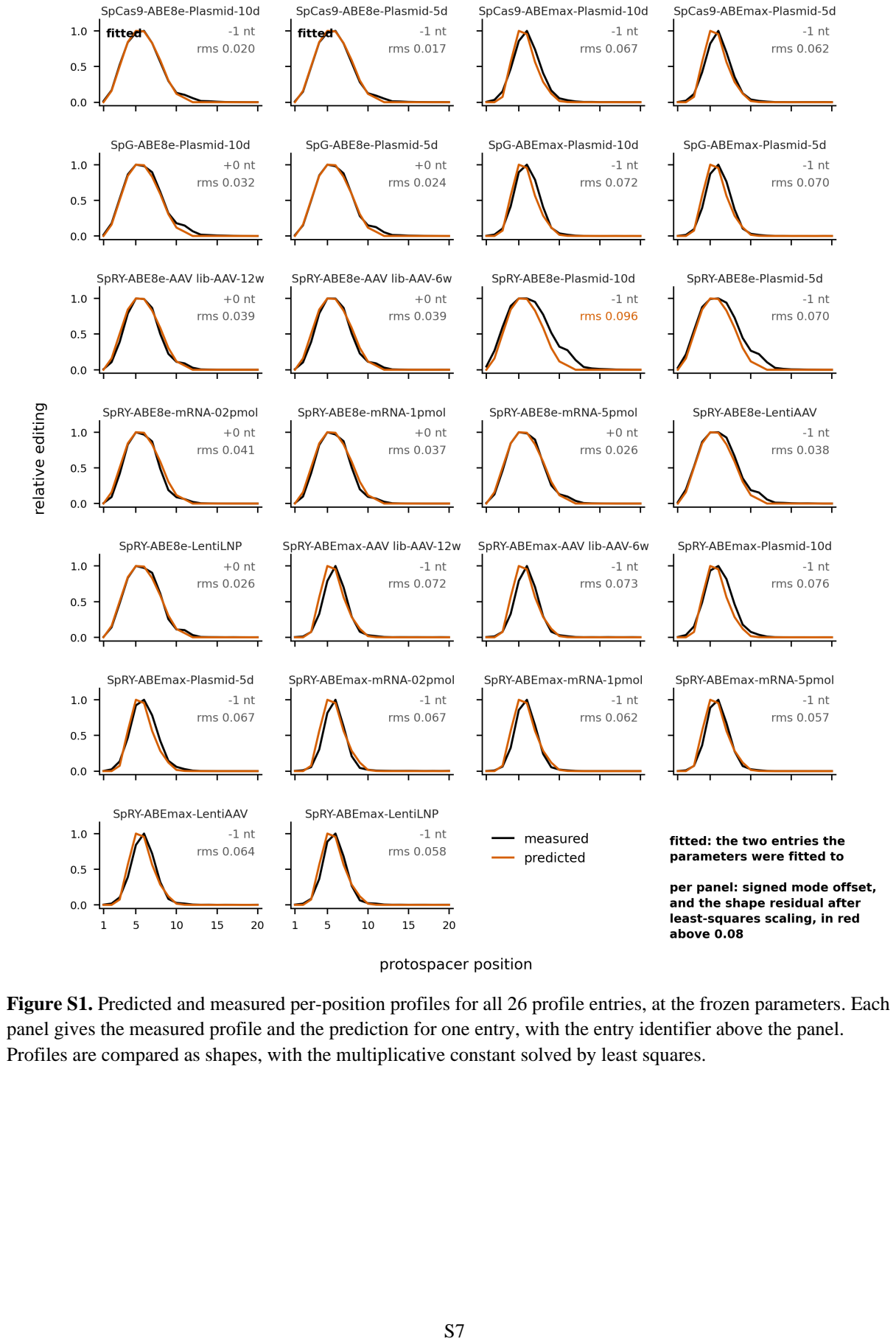

Figure S1. Predicted and measured per-position profiles for all 26 profile entries, at the frozen parameters. Each panel gives the measured profile and the prediction for one entry, with the entry identifier above the panel. Profiles are compared as shapes, with the multiplicative constant solved by least squares.

**Table S1.** Corpus of 50 architecture entries with per-entry provenance.

| Entry | Source | Editor | Ortholog | Anchor site | Linker res. | Linker class | Effector | Label | Readout | Context | Reported window | Panel or table |
| --- | --- | --- | --- | --- | --- | --- | --- | --- | --- | --- | --- | --- |
| kissling-SpRY-ABE8e-HEK-Plasmid-5d | kissling2025 | SpRY-ABE8e | SpRY | n_terminal | 32 | xten | tada8e | effector_varied | amplicon_sequencing | HEK-Plasmid_5d |  | Additional file 4, sheet HEK-Plasmid_5d_SpRY-ABE8e |
| kissling-SpRY-ABE8e-HEK-Plasmid-10d | kissling2025 | SpRY-ABE8e | SpRY | n_terminal | 32 | xten | tada8e | effector_varied | amplicon_sequencing | HEK-Plasmid_10d |  | Additional file 4, sheet HEK-Plasmid_10d_SpRY-ABE8e |
| kissling-SpRY-ABE8e-HEK-mRNA-02pmol | kissling2025 | SpRY-ABE8e | SpRY | n_terminal | 32 | xten | tada8e | effector_varied | amplicon_sequencing | HEK-mRNA_02pmol |  | Additional file 4, sheet HEK-mRNA_02pmol_SpRY-ABE8e |
| kissling-SpRY-ABE8e-HEK-mRNA-1pmol | kissling2025 | SpRY-ABE8e | SpRY | n_terminal | 32 | xten | tada8e | effector_varied | amplicon_sequencing | HEK-mRNA_1pmol |  | Additional file 4, sheet HEK-mRNA_1pmol_SpRY-ABE8e |
| kissling-SpRY-ABE8e-HEK-mRNA-5pmol | kissling2025 | SpRY-ABE8e | SpRY | n_terminal | 32 | xten | tada8e | effector_varied | amplicon_sequencing | HEK-mRNA_5pmol |  | Additional file 4, sheet HEK-mRNA_5pmol_SpRY-ABE8e |
| kissling-SpRY-ABE8e-AAVlibrary-AAV-6w | kissling2025 | SpRY-ABE8e | SpRY | n_terminal | 32 | xten | tada8e | effector_varied | amplicon_sequencing | AAVlibrary-AAV_6w |  | Additional file 4, sheet AAVlibrary-AAV_6w_SpRY-ABE8e |
| kissling-SpRY-ABE8e-AAVlibrary-AAV-12w | kissling2025 | SpRY-ABE8e | SpRY | n_terminal | 32 | xten | tada8e | effector_varied | amplicon_sequencing | AAVlibrary-AAV_12w |  | Additional file 4, sheet AAVlibrary-AAV_12w_SpRY-ABE8e |
| kissling-SpRY-ABE8e-LentiAAV | kissling2025 | SpRY-ABE8e | SpRY | n_terminal | 32 | xten | tada8e | effector_varied | amplicon_sequencing | LentiAAV |  | Additional file 4, sheet LentiAAV_SpRY-ABE8e |
| kissling-SpRY-ABE8e-LentiLNP | kissling2025 | SpRY-ABE8e | SpRY | n_terminal | 32 | xten | tada8e | effector_varied | amplicon_sequencing | LentiLNP |  | Additional file 4, sheet LentiLNP_SpRY-ABE8e |
| kissling-SpRY-ABEmax-HEK-Plasmid-5d | kissling2025 | SpRY-ABEmax | SpRY | n_terminal | 32 | xten | abemax | scaffold_perturbed | amplicon_sequencing | HEK-Plasmid_5d |  | Additional file 4, sheet HEK-Plasmid_5d_SpRY-ABEmax |
| kissling-SpRY-ABEmax-HEK-Plasmid-10d | kissling2025 | SpRY-ABEmax | SpRY | n_terminal | 32 | xten | abemax | scaffold_perturbed | amplicon_sequencing | HEK-Plasmid_10d |  | Additional file 4, sheet HEK-Plasmid_10d_SpRY-ABEmax |
| kissling-SpRY-ABEmax-HEK-mRNA-02pmol | kissling2025 | SpRY-ABEmax | SpRY | n_terminal | 32 | xten | abemax | scaffold_perturbed | amplicon_sequencing | HEK-mRNA_02pmol |  | Additional file 4, sheet HEK-mRNA_02pmol_SpRY-ABEmax |
| kissling-SpRY-ABEmax-HEK-mRNA-1pmol | kissling2025 | SpRY-ABEmax | SpRY | n_terminal | 32 | xten | abemax | scaffold_perturbed | amplicon_sequencing | HEK-mRNA_1pmol |  | Additional file 4, sheet HEK-mRNA_1pmol_SpRY-ABEmax |
| kissling-SpRY-ABEmax-HEK-mRNA-5pmol | kissling2025 | SpRY-ABEmax | SpRY | n_terminal | 32 | xten | abemax | scaffold_perturbed | amplicon_sequencing | HEK-mRNA_5pmol |  | Additional file 4, sheet HEK-mRNA_5pmol_SpRY-ABEmax |
| kissling-SpRY-ABEmax-AAVlibrary-AAV-6w | kissling2025 | SpRY-ABEmax | SpRY | n_terminal | 32 | xten | abemax | scaffold_perturbed | amplicon_sequencing | AAVlibrary-AAV_6w |  | Additional file 4, sheet AAVlibrary-AAV_6w_SpRY-ABEmax |
| kissling-SpRY-ABEmax-AAVlibrary-AAV-12w | kissling2025 | SpRY-ABEmax | SpRY | n_terminal | 32 | xten | abemax | scaffold_perturbed | amplicon_sequencing | AAVlibrary-AAV_12w |  | Additional file 4, sheet AAVlibrary-AAV_12w_SpRY-ABEmax |
| kissling-SpRY-ABEmax-LentiAAV | kissling2025 | SpRY-ABEmax | SpRY | n_terminal | 32 | xten | abemax | scaffold_perturbed | amplicon_sequencing | LentiAAV |  | Additional file 4, sheet LentiAAV_SpRY-ABEmax |
| kissling-SpRY-ABEmax-LentiLNP | kissling2025 | SpRY-ABEmax | SpRY | n_terminal | 32 | xten | abemax | scaffold_perturbed | amplicon_sequencing | LentiLNP |  | Additional file 4, sheet LentiLNP_SpRY-ABEmax |
| kissling-SpG-ABE8e-HEK-Plasmid-5d | kissling2025 | SpG-ABE8e | SpG | n_terminal | 32 | xten | tada8e | effector_varied | amplicon_sequencing | HEK-Plasmid_5d |  | Additional file 4, sheet HEK-Plasmid_5d_SpG-ABE8e |
| kissling-SpG-ABE8e-HEK-Plasmid-10d | kissling2025 | SpG-ABE8e | SpG | n_terminal | 32 | xten | tada8e | effector_varied | amplicon_sequencing | HEK-Plasmid_10d |  | Additional file 4, sheet HEK-Plasmid_10d_SpG-ABE8e |
| kissling-SpG-ABEmax-HEK-Plasmid-5d | kissling2025 | SpG-ABEmax | SpG | n_terminal | 32 | xten | abemax | scaffold_perturbed | amplicon_sequencing | HEK-Plasmid_5d |  | Additional file 4, sheet HEK-Plasmid_5d_SpG-ABEmax |
| kissling-SpG-ABEmax-HEK-Plasmid-10d | kissling2025 | SpG-ABEmax | SpG | n_terminal | 32 | xten | abemax | scaffold_perturbed | amplicon_sequencing | HEK-Plasmid_10d |  | Additional file 4, sheet HEK-Plasmid_10d_SpG-ABEmax |
| kissling-SpCas9-ABE8e-HEK-Plasmid-5d | kissling2025 | SpCas9-ABE8e | SpCas9 | n_terminal | 32 | xten | tada8e | effector_varied | amplicon_sequencing | HEK-Plasmid_5d |  | Additional file 4, sheet HEK-Plasmid_5d_SpCas9-ABE8e |
| kissling-SpCas9-ABE8e-HEK-Plasmid-10d | kissling2025 | SpCas9-ABE8e | SpCas9 | n_terminal | 32 | xten | tada8e | effector_varied | amplicon_sequencing | HEK-Plasmid_10d |  | Additional file 4, sheet HEK-Plasmid_10d_SpCas9-ABE8e |
| kissling-SpCas9-ABEmax-HEK-Plasmid-5d | kissling2025 | SpCas9-ABEmax | SpCas9 | n_terminal | 32 | xten | abemax | none | amplicon_sequencing | HEK-Plasmid_5d |  | Additional file 4, sheet HEK-Plasmid_5d_SpCas9-ABEmax |
| kissling-SpCas9-ABEmax-HEK-Plasmid-10d | kissling2025 | SpCas9-ABEmax | SpCas9 | n_terminal | 32 | xten | abemax | none | amplicon_sequencing | HEK-Plasmid_10d |  | Additional file 4, sheet HEK-Plasmid_10d_SpCas9-ABEmax |
| hnhx-ABEmax7.10 | hnhx2021 | ABEmax7.10 | SpCas9 | n_terminal | 32 | xten | abemax | none | amplicon_sequencing | HEK293T | 4-8 | Discussion, window quoted as positions 4 to 8 |
| hnhx-ABEmax7.10-dHNH | hnhx2021 | ABEmax7.10 dHNH | SpCas9 | n_terminal | 32 | xten | abemax | scaffold_perturbed | amplicon_sequencing | HEK293T |  | Figure 1E, editing gained at positions 12 and 14 |
| hnhx-ABEmax7.10-inserted | hnhx2021 | HNHx-ABEmax7.10 | SpCas9 | hnh_amino | 3 | gs | tada_inserted | scaffold_perturbed | amplicon_sequencing | HEK293T | 7-14 | Figure 3A, window quoted as positions 7 to 14 |
| hnhx-ABE8e-inserted | hnhx2021 | HNHx-ABE8e | SpCas9 | hnh_amino | 3 | gs | tada_inserted | scaffold_perturbed | amplicon_sequencing | HEK293T |  | Figure 3B, shift retained and the window broadened |
| komor-linker-3 | komor2016 | rAPOBEC1-dCas9 | SpCas9 | n_terminal | 3 | gs | apobec1 | tether_varied | in_vitro_deamination | purified protein on dsDNA |  | Extended Data Figure 1c, window about three nucleotides wide |
| komor-linker-16 | komor2016 | rAPOBEC1-dCas9 | SpCas9 | n_terminal | 16 | xten | apobec1 | tether_varied | in_vitro_deamination | purified protein on dsDNA | 4-8 | Extended Data Figure 1e and the text, positions 4 to 8 |
| komor-linker-21 | komor2016 | rAPOBEC1-dCas9 | SpCas9 | n_terminal | 21 | gs | apobec1 | tether_varied | in_vitro_deamination | purified protein on dsDNA |  | Extended Data Figure 1f, window about six nucleotides wide |
| bepigs-BE3 | bepigs2019 | BE3 | SpCas9 | n_terminal | 16 | xten | apobec1 | none | sanger_deconvolution | HEK293T | 4-8 | Figure 1g, window quoted as positions 4 to 8 |
| bepigs-pi-16 | bepigs2019 | BE-PIGS | SpCas9 | pi_amino | 16 | xten | apobec1_inserted | scaffold_perturbed | sanger_deconvolution | HEK293T | 4-14 | Figure 1g, positions 4 to 14 with the maximum around 7 to 13 |
| bepigs-pi-8 | bepigs2019 | BE-PIGS N8 | SpCas9 | pi_amino | 8 | xten | apobec1_inserted | scaffold_perturbed | sanger_deconvolution | HEK293T |  | Figure S3, efficiency fell and the width did not obviously change |
| bepigs-pi-3 | bepigs2019 | BE-PIGS N3 | SpCas9 | pi_amino | 3 | xten | apobec1_inserted | scaffold_perturbed | sanger_deconvolution | HEK293T |  | Figure S3, efficiency fell and the width did not obviously change |
| bepigs-ruvc-16 | bepigs2019 | BE-RuvCGE | SpCas9 | ruvc_amino | 16 | xten | apobec1_inserted | scaffold_perturbed | sanger_deconvolution | HEK293T |  | Figure 1c and 1d, editing about half that of the PAM-interacting inlay |
| huangcp-ABEmax | huangcp | ABEmax | SpCas9 | n_terminal | 32 | xten | abemax | none | amplicon_sequencing | HEK293T | 4-7 | Figure 2f, canonical window quoted as positions 4 to 7 with the maximum at 5 or 6 |
| huangcp-CP1012-ABEmax | huangcp | CP1012-ABEmax | SpCas9 | cp1012 | 32 | xten | abemax | scaffold_perturbed | amplicon_sequencing | HEK293T | 4-12 | Figure 2f, window 4 to 12; the maximum moves to 6 or 7 |
| huangcp-CP1028-ABEmax | huangcp | CP1028-ABEmax | SpCas9 | cp1028 | 32 | xten | abemax | scaffold_perturbed | amplicon_sequencing | HEK293T | 4-12 | Figure 2f, window 4 to 12 |
| huangcp-CP1041-ABEmax | huangcp | CP1041-ABEmax | SpCas9 | cp1041 | 32 | xten | abemax | scaffold_perturbed | amplicon_sequencing | HEK293T | 4-12 | Figure 2f, window 4 to 12; the broadest, with editing at 14 |
| huangcp-CP1249-ABEmax | huangcp | CP1249-ABEmax | SpCas9 | cp1249 | 32 | xten | abemax | scaffold_perturbed | amplicon_sequencing | HEK293T | 4-12 | Figure 2f, window 4 to 12 |
| tan2019-xten-16 | tan2019 | BE3 | SpCas9 | n_terminal | 16 | xten | apobec1 | none | functional_selection | yeast CAN1 selection |  | Figure 1b, editing at every cytosine across a nine nucleotide window |
| tan2019-pap-3 | tan2019 | BE3 PAP | SpCas9 | n_terminal | 3 | polyproline | apobec1 | tether_varied | functional_selection | yeast CAN1 selection |  | Figure 1b, editing abolished almost completely |
| tan2019-papap-5 | tan2019 | BE3 PAPAP | SpCas9 | n_terminal | 5 | polyproline | apobec1 | tether_varied | functional_selection | yeast CAN1 selection | 5-7 | Supplementary Figure 5, editing mainly at positions 14 to 16 from the PAM |
| tan2019-papapap-7 | tan2019 | BE3 PAPAPAP | SpCas9 | n_terminal | 7 | polyproline | apobec1 | tether_varied | functional_selection | yeast CAN1 selection | 5-6 | Figure 1b, editing largely restricted to positions 15 and 16 from the PAM |
| communbiol-dCas12a-A3A | communbiol2022 | dCas12a-A3A | Cas12a | n_terminal | 16 | xten | ha3a | none | amplicon_sequencing | HEK293T | 6-15 | Supplementary Figure 3, window quoted as positions 6 to 15; the amino-terminal arrangement and the 16-residue XTEN are read from the deposited sequence of Addgene 193640, not inferred |
| communbiol-dCas12a-A3A-Y130F | communbiol2022 | dCas12a-A3A-Y130F | Cas12a | n_terminal | 16 | xten | ha3a | effector_varied | amplicon_sequencing | HEK293T | 6-12 | Table 1, window quoted as positions 6 to 12; the amino-terminal arrangement and the 16-residue XTEN are read from the deposited sequence of Addgene 193640, not inferred |
| communbiol-dCas12a-ABE8e | communbiol2022 | dCas12a-ABE8e | Cas12a | n_terminal | 16 | xten | tada8e | effector_varied | amplicon_sequencing | HEK293T | 8-12 | Supplementary Figure 8, window quoted as positions 8 to 12; the amino-terminal arrangement and the 16-residue XTEN are read from the deposited sequence of Addgene 193640, not inferred |

Anchor site, linker composition and residue count, effector, readout class and the panel or table from which each value was read. Entries read out by amplicon sequencing contribute to fitting and to the width analyses; the others are used for evaluation only. The columns shown here are the ones the text refers to; the complete twenty-one-column record, including spacer length, protospacer numbering convention, structure identifier, scaffold variant, return-linker site and residue count, and reported window width, is on the Table S1 sheet of SI\_data.xls.

**Table S2.** Source register: 28 entries covering 29 studies, with role, status and, for each exclusion, the reason.

| Key | Citation | DOI | Role | Tether variation | Readout | Status | Notes |
| --- | --- | --- | --- | --- | --- | --- | --- |
| komor2016 | Komor AC, Kim YB, Packer MS, Zuris JA, Liu DR. Programmable editing of a target base in genomic DNA without double-stranded DNA cleavage. *Nature* 2016;533:420-424 | 10.1038/nature17946 | architect<br>ure | linker_length | in_vitro_deamination | confirmed | Purified rAPOBEC1-dCas9 on double-stranded DNA with linkers stated as GGS, (GGS)3, a 16-residue XTEN and (GGS)7, being 3, 9, 16 and 21 residues. Window width read from the gel panels and quoted in the text. The only source that states linker length in residues, and the width claims rest on it |
| koblan2018 | Koblan LW, Doman JL, Wilson C, Levy JM, Tay T, Newby GA, Maianti JP, Raguram A, Liu DR. Improving cytidine and adenine base editors by expression optimization and ancestral reconstruction. *Nat Biotechnol* 2018;36:843-846 | 10.1038/nbt.4172 | architect<br>ure | none | amplicon_sequencing | no_tether_variation | The improvements are nuclear localisation and codon usage; the linkers are unchanged and the editing window is reported unchanged. A specificity control rather than an architecture source |
| tan2019 | Tan J, Zhang F, Karcher D, Bock R. Engineering of high-precision base editors for site-specific single nucleotide replacement. *Nat Commun* 2019;10:439 | 10.1038/s41467-018-08034-8 | architect<br>ure | rigidity_and_terminus | functional_selection | confirmed | Ten rigid proline-containing linkers against the 16-residue XTEN. Three residues and no linker abolish editing, five to seven narrow the window to positions 14 to 16 from the PAM, and longer ones widen it again. Positions are quoted relative to the PAM and converted on the reading that the last protospacer base is -1 |
| tan2020 | Tan J, Zhang F, Karcher D, Bock R. Expanding the genome-targeting scope and the site selectivity of high-precision base editors. *Nat Commun* 2020;11:629 | 10.1038/s41467-020-14465-z | architect<br>ure | deaminase_truncation | functional_selection | effector_geometry_mismatching | Varies the carboxy-terminal tail of CDA1 by 13 to 20 residues, which is tether length expressed through the effector rather than through the linker. No structure is assigned for CDA1, so no offsets or body radius can be measured and no entry can be built |
| bepigs2019 | Wang Y, Zhou L, Liu N, Yao S. BE-PIGS: a base-editing tool with deaminases inlaid into Cas9 PI domain significantly expanded the editing scope. *Signal Transduct Target Ther* 2019;4:36 | 10.1038/s41392-019-0072-7 | architect<br>ure | inlay_and_linker_length | sanger_deconvolution | confirmed | APOBEC1 inlaid between G1054 and E1055 and between G1246 and S1247, with an amino-terminal linker of 16, 8 or 3 residues and a return linker of 32. The PAM-interacting inlay edits positions 4 to 14 |
| hnhx2021 | Villiger L, Schmidheini L, Mathis N, Rothgangl T, Marquart K, Schwank G. Replacing the SpCas9 HNH domain by deaminases generates compact base editors with an alternative targeting scope. *Mol Ther Nucleic Acids* 2021;26:502-510 | 10.1016/j.omtn.2021.08.025 | architect<br>ure | anchor_transfer_and_hnh_removal | amplicon_sequencing | confirmed | HNH taken as residues 775 to 908. The insertion joins S793 to the TadA amino terminus through GGS and the TadA carboxy terminus to R919 through SGG. Windows are quoted in the text and per-position values appear only in figures |
| huangcp | Huang TP, Zhao KT, Miller SM, Gaudelli NM, Oakes BL, Fellmann C, Savage DF, Liu DR. Circularly permuted and PAM-modified Cas9 variants broaden the targeting scope of base editors. *Nat Biotechnol* 2019;37:626-631 | 10.1038/s41587-019-0134-y | architect<br>ure | anchor_reorganisation | amplicon_sequencing | confirmed | Five permutants, CP1012, CP1028, CP1041, CP1249 and CP1300, the number naming the new amino terminus. ABEmax edits positions 4 to 7 and the permutants 4 to 12, with CP1041 the broadest |
| nme2inlay2023 | Bamidele N, Zhang H, Dong X, Cheng H, Gaston N, Feinzig H, Cao H, Kelly K, Watts JK, Xie J, Gao G, Sontheimer EJ. Domain-inlaid Nme2Cas9 adenine base editors with improved activity and targeting scope. *Nat Commun* 2024;15:1697 | 10.1038/s41467-024-45763-5 | architect<br>ure | domain_insertion | amplicon_sequencing | insertion_residues_unavailable | Eight insertion sites with TadA8e between two 20-residue linkers; i1 edits PAM-distally and i7 and i8 PAM-proximally. The residue numbers appear only in a figure panel, and the structure used is Nme1Cas9 rather than Nme2Cas9, so no entry can be built |
| li2018 | Li X, Wang Y, Liu Y, Yang B, Wang X, Wei J, Lu Z, Zhang Y, Wu J, Huang X, Yang L, Chen J. Base editing with a Cpf1-cytidine deaminase fusion. *Nat Biotechnol* 2018;36:324-327 | 10.1038/nbt.4102 | architect<br>ure | cas12a_architecture | amplicon_sequencing | not_open_access | Not in the open-access set. Its construct map, Addgene 107685, is the arrangement the later Cas12a base editors inherit, and the same arrangement appears in the plasmid Chen and colleagues deposited |
| plantcomm2023 | Cheng Y, Zhang Y, Li G, Fang H, Sretenovic S, Fan A, Li J, Xu J, Que Q, Qi Y. CRISPR-Cas12a base editors confer efficient multiplexed genome editing in rice. *Plant Commun* 2023;4:100601 | 10.1016/j.xplc.2023.100601 | architect<br>ure | linker_length | plant_system | linker_lengths_unavailable | dLbCas12a-D156R ABE edits A8 to A11 and the CBE prefers C10, but the seven connector sequences appear only in a figure panel, so no entry can record a contour length |
| communbiol 2022 | Chen F, Lian M, Ma B, Gou S, Luo X, Yang K, Shi H, Xie J, Ge W, Ouyang Z, Lai C, Li N, Zhang Q, Jin Q, Liang Y, Chen T, Wang J, Zhao X, Li L, Yu M, Ye Y, Wang K, Wu H, Lai L. Multiplexed base editing through Cas12a variant-mediated cytosine and adenine base editors. Commun Biol 2022;5:1163 | 10.1038/s42003-022-04152-8 | architecture | cas12a_scaffold_and_deaminase | amplicon_sequencing | confirmed | Windows in Cas12a numbering: hA3A 6 to 15, hA3A-Y130F 6 to 12, TadA8e 8 to 12. The paper states neither the linker nor the terminus. Both were taken from the deposited sequence of the authors' plasmid, Addgene 193640: human APOBEC3A at 409 to 1005 with phenylalanine at residue 130, then SGSETPGTSESATPES, then LbCas12a. An amino-terminal deaminase on a 16-residue XTEN |
| abenw1_2026 | Valdez I, O'Connor I, Patel D, Gierer K, Harrington J, Ellis E, Caponetti SA, Sebra RP, Valley HC, Coote K, Mense M, Marro SG, Jiang T. A streamlined base editor engineering strategy to reduce bystander editing. Nat Commun 2025;16:8115 | 10.1038/s41467-025-63609-6 | architecture | none | amplicon_sequencing | outside_the_model | TadA-NW1 carries a naturally occurring oligonucleotide binding module in its own deaminase active centre, so the window narrows from the ten nucleotides of TadA-8e-derived ABEs to four through the deaminase binding the substrate rather than through the tether. Outside what a tether model represents |
| shi2026 | Shi Y, Yuan Y, Qin L, Zhou F, Wu G, Li B, Yao P, Shi M, Ma L, Wang Y, Zhang Y, Wang C, Wang X, Huang B, Chen J, Xiang Z, Lin Q, Huang J. Optimizing linker length of base editors for precise crop breeding and gene therapy. J Genet Genomics 2026;53:1125-1137 | 10.1016/j.jgg.2026.02.021 | architecture | linker_length | amplicon_sequencing | not_open_access | Not in the open-access set at PubMed Central |
| kuang2021 | Kuang J, Lyu Q, Wang J, Cui Y, Zhao J. Advances in base editing with an emphasis on an AAV-based strategy. Methods 2021;194:56-64 | 10.1016/j.ymeth.2021.03.015 | architecture | linker_length | amplicon_sequencing | not_open_access | Cited as prior linker work by Shi and colleagues. A review that surveys linker length among other optimisations rather than reporting an architecture series of its own, and the full text is not open access |
| zhao2022 | Zhao D, Qian Y, Li J, Li Z, Lai L. Highly efficient A-to-G base editing by ABE8.17 in rabbits. Mol Ther Nucleic Acids 2022;27:1156-1163 | 10.1016/j.omtn.2022.01.019 | architecture | linker_length | amplicon_sequencing | outside_the_model | Cited as prior linker work by Shi and colleagues. The tether variation is a deletion: ABE8.17 uses a 32-residue (SGGS)2-XTEN-(SGGS)2 linker and ABE8.17-NL removes it entirely, narrowing the reported window to 2 to 4 nucleotides. A zero-residue linker gives the model no chain to grow, so the pair cannot be scored as a length contrast, and the 32-residue parent repeats an amino-terminal XTEN-32 architecture already in the corpus |
| iscb_family | Han D, Xiao Q, Wang Y, Zhang H, Dong X, Li G, et al. Development of miniature base editors using engineered IscB nickase. Nat Methods 2023;20:1029-1036; and Han L, Hu Y, Mo Q, Yang H, Gu F, Bai F, et al. Engineering miniature IscB nickase for robust base editing with broad targeting range. Nat Chem Biol 2024;20:1629-1639 | 10.1038/s41592-023-01898-9; 10.1038/s41589-024-01670-w | architecture | compact_scaffold | amplicon_sequencing | no_structure | Two studies held as one register row: miBE and InScB from the first, IminiBE and SIminiBE from the second. Minimal tether variation, and no R-loop structure is assigned for IscB, so nothing can be placed in a frame |
| kim2024 | Kim N, Choi S, Kim S, Song M, Seo JH, Min S, Park J, Cho SR, Kim HH. Deep learning models to predict the editing efficiencies and outcomes of diverse base editors. Nat Biotechnol 2024;42:484-497 | 10.1038/s41587-023-01792-x | profile | none | amplicon_sequencing | confirmed | Windows, outcomes and motifs for seven base editors and nine Cas9 variants in one assay |
| kissling2025 | Kissling L, Mollaysa A, Janjuha S, Mathis N, Marquart KF, Weber Y, Moon WJ, Lin PJC, Fan SHY, Muramatsu H, Vadovics M, Allam A, Pardi N, Tam YK, Krauthammer M, Schwank G. Predicting adenine base editing efficiencies in different cellular contexts by deep learning. Genome Biol 2025;26:115 | 10.1186/s13059-025-03586-7 | profile | none | amplicon_sequencing | confirmed | Per-position efficiency for six architectures, in vitro and in vivo, curated from Additional file 4. The paper names ABEmax and ABE8e without stating a linker length, so the 32 residues those entries carry come from the construct architecture rather than from this source |
| marquart2021 | Marquart KF, Allam A, Janjuha S, Sintsova A, Villiger L, Frey N, Krauthammer M, Schwank G. Predicting base editing outcomes with an attention-based deep learning algorithm trained on high-throughput target library screens. Nat Commun 2021;12:5114 | 10.1038/s41467-021-25375-z | profile | none | amplicon_sequencing | confirmed | Four architectures against one library of 28,000 targets |
| arbab2020 | Arbab M, Shen MW, Wilson C, Matuszek ■, Cassa CA, Liu DR. Determinants of Base Editing Outcomes from Target Library Analysis and Machine Learning. Cell 2020;182:463-480.e30 | 10.1016/j.cell.2020.05.037 | profile | none | amplicon_sequencing | confirmed | Eleven editors across more than 38,000 sites |
| pallaseni2022 | Pallaseni A, Peets EM, Koeppel J, Weller J, Vanderstichele T, Ho UL, Crepaldi L, van Leeuwen J, Allen F, Parts L. Predicting base editing outcomes using position-specific sequence determinants. *Nucleic Acids Res* 2022;50:3551-3564 | 10.1093/nar/gkac161 | profile | none | amplicon_sequencing | confirmed | Position-specific determinants in two cell lines |
| zhou2024 | Zhou X, Gao J, Luo L, Huang C, Wu J, Wang X. Comprehensive evaluation and prediction of editing outcomes for near-PAMless adenine and cytosine base editors. *Commun Biol* 2024;7:1389 | 10.1038/s42003-024-07078-5 | profile | none | amplicon_sequencing | confirmed | Near-PAMless ABE and CBE |
| bedatahive2024 | Schneider L, Minary P. Be-dataHIVE: a base editing database. *BMC Bioinformatics* 2024;25:330 | 10.1186/s12859-024-05898-0 | profile | none | amplicon_sequencing | confirmed | Aggregated schema |
| feola2024 | Feola M, Pulicani S, Tkach D, Boyne A, Hong R, Mayer L, Duclert A, Duchateau P, Juillerat A. Comprehensive analysis of the editing window of C-to-T TALE base editors. *Sci Rep* 2024;14:12870 | 10.1038/s41598-024-63203-8 | external | tale_linker | amplicon_sequencing | confirmed | Three connector scaffolds between the TALE array and the split deaminase: C40 at about 43 residues, C11 at 20 and C0 at 3. Both halves carry a scaffold, so the connector is varied on both sides and the check stays qualitative. C0 does not edit at any spacer length |
| xiang2025 | Xiang J, Xu W, Wu J, Luo Y, Liu C, Hou Y, Chen J, Yang B. Structural insights into DdCBE in action enable high-precision mitochondrial DNA editing. *Mol Cell* 2025;85:3357-3372.e9 | 10.1016/j.molcel.2025.08.016 | external | none | amplicon_sequencing | confirmed | Cryo-electron microscopy of DdCBE, with the WinPred window predictor |
| mi2025 | Mi L, Li YX, Lv X, Wan ZL, Liu X, Zhang K, Li H, Yao Y, Zhang L, Xu Z, Zhuang X, Ji K, Jiang M, Wang Y, Lu P. Computational design of a high-precision mitochondrial DNA cytosine base editor. *Nat Struct Mol Biol* 2025;32:2575-2586 | 10.1038/s41594-025-01714-2 | external | none | amplicon_sequencing | confirmed | Designed rigid TALE-deaminase interface, named TALE-oriented deaminase |
| evers2006 | Evers TH, van Dongen EMWM, Faesen AC, Meijer EW, Merkx M. Quantitative understanding of the energy transfer between fluorescent proteins connected via flexible peptide linkers. *Biochemistry* 2006;45:13183-13192 | 10.1021/bi061288t | external | none | fret | confirmed | Worm-like-chain fit to FRET between fluorescent domains joined by GlyGlySer repeats gives a persistence length of 4.5 angstrom. The primary source for the gs composition class |
| vanrosmalen2017 | van Rosmalen M, Krom M, Merkx M. Tuning the flexibility of glycine-serine linkers to allow rational design of multidomain proteins. *Biochemistry* 2017;56:6565-6574 | 10.1021/acs.biochem.7b00902 | external | none | fret | confirmed | Worm-like-chain fits across a glycine series give 4.5 angstrom at 33 percent glycine, 4.8 at 17 percent and 6.2 with none, reproducing Evers at 4.5. These bracket the xten class, for which no direct measurement is available, so that class is banded over 4.5 to 6.4 rather than pinned |

**Table S3.** Fitted parameters and loss across the five fitting subsets, with the corpus hit rate under each parameter set.

| Fitting subset | n fitted | Capture radius (Å) | Strand persistence (Å) | Link steepness | Loss | Entries scored | Within one nucleotide |
| --- | --- | --- | --- | --- | --- | --- | --- |
| training split, SpCas9-ABE8e plasmid | 2 | 22.0 | 12.0 | 740.6 | 0.015 | 39 | 0.85 |
| every SpCas9 terminal fusion with a profile | 4 | 25.0 | 40.0 | 90.5 | 0.095 | 41 | 0.83 |
| ABE8e only | 13 | 21.0 | 10.0 | 1492.5 | 0.468 | 39 | 0.85 |
| ABEmax only | 13 | 22.0 | 8.0 | 367.5 | 0.371 | 39 | 0.82 |
| every profile entry | 26 | 21.0 | 8.0 | 1492.5 | 0.975 | 39 | 0.82 |

**Table S4A.** Predicted mode at five sampling seeds, by entry.

| Entry | Ortholog | Observed | seed 0 | seed 1 | seed 2 | seed 3 | seed 4 | Spread |
| --- | --- | --- | --- | --- | --- | --- | --- | --- |
| kissling-SpRY-ABE8e-HEK-Plasmid-5d | SpRY | 6 | 5 | 5 | 5 | 6 | 5 | 1 |
| kissling-SpRY-ABE8e-HEK-Plasmid-10d | SpRY | 6 | 5 | 5 | 5 | 6 | 5 | 1 |
| kissling-SpRY-ABE8e-HEK-mRNA-02pmol | SpRY | 5 | 5 | 5 | 5 | 6 | 5 | 1 |
| kissling-SpRY-ABE8e-HEK-mRNA-1pmol | SpRY | 5 | 5 | 5 | 5 | 6 | 5 | 1 |
| kissling-SpRY-ABE8e-HEK-mRNA-5pmol | SpRY | 5 | 5 | 5 | 5 | 6 | 5 | 1 |
| kissling-SpRY-ABE8e-AAVlibrary-AAV-6w | SpRY | 5 | 5 | 5 | 5 | 6 | 5 | 1 |
| kissling-SpRY-ABE8e-AAVlibrary-AAV-12w | SpRY | 5 | 5 | 5 | 5 | 6 | 5 | 1 |
| kissling-SpRY-ABE8e-LentiAAV | SpRY | 6 | 5 | 5 | 5 | 6 | 5 | 1 |
| kissling-SpRY-ABE8e-LentiLNP | SpRY | 5 | 5 | 5 | 5 | 6 | 5 | 1 |
| kissling-SpRY-ABEmax-HEK-Plasmid-5d | SpRY | 6 | 6 | 5 | 6 | 6 | 6 | 1 |
| kissling-SpRY-ABEmax-HEK-Plasmid-10d | SpRY | 6 | 6 | 5 | 6 | 6 | 6 | 1 |
| kissling-SpRY-ABEmax-HEK-mRNA-02pmol | SpRY | 6 | 6 | 5 | 6 | 6 | 6 | 1 |
| kissling-SpRY-ABEmax-HEK-mRNA-1pmol | SpRY | 6 | 6 | 5 | 6 | 6 | 6 | 1 |
| kissling-SpRY-ABEmax-HEK-mRNA-5pmol | SpRY | 6 | 6 | 5 | 6 | 6 | 6 | 1 |
| kissling-SpRY-ABEmax-AAVlibrary-AAV-6w | SpRY | 6 | 6 | 5 | 6 | 6 | 6 | 1 |
| kissling-SpRY-ABEmax-AAVlibrary-AAV-12w | SpRY | 6 | 6 | 5 | 6 | 6 | 6 | 1 |
| kissling-SpRY-ABEmax-LentiAAV | SpRY | 6 | 6 | 5 | 6 | 6 | 6 | 1 |
| kissling-SpRY-ABEmax-LentiLNP | SpRY | 6 | 6 | 5 | 6 | 6 | 6 | 1 |
| kissling-SpG-ABE8e-HEK-Plasmid-5d | SpG | 5 | 5 | 5 | 5 | 6 | 5 | 1 |
| kissling-SpG-ABE8e-HEK-Plasmid-10d | SpG | 5 | 5 | 5 | 5 | 6 | 5 | 1 |
| kissling-SpG-ABEmax-HEK-Plasmid-5d | SpG | 6 | 6 | 5 | 6 | 6 | 6 | 1 |
| kissling-SpG-ABEmax-HEK-Plasmid-10d | SpG | 6 | 6 | 5 | 6 | 6 | 6 | 1 |
| kissling-SpCas9-ABE8e-HEK-Plasmid-5d | SpCas9 | 6 | 5 | 5 | 5 | 6 | 5 | 1 |
| kissling-SpCas9-ABE8e-HEK-Plasmid-10d | SpCas9 | 6 | 5 | 5 | 5 | 6 | 5 | 1 |
| kissling-SpCas9-ABEmax-HEK-Plasmid-5d | SpCas9 | 6 | 6 | 5 | 6 | 6 | 6 | 1 |
| kissling-SpCas9-ABEmax-HEK-Plasmid-10d | SpCas9 | 6 | 6 | 5 | 6 | 6 | 6 | 1 |
| hnhx-ABEmax7.10 | SpCas9 | 6 | 6 | 5 | 6 | 6 | 6 | 1 |
| hnhx-ABEmax7.10-inserted | SpCas9 | 10 | no window | 15 | no window | 15 | no window | 0 |
| komor-linker-16 | SpCas9 | 6 | 5 | 5 | 5 | 5 | 5 | 0 |
| bepigs-BE3 | SpCas9 | 6 | 5 | 5 | 5 | 5 | 5 | 0 |
| bepigs-pi-16 | SpCas9 | 9 | 11 | 11 | 11 | 11 | 11 | 0 |
| huangcp-ABEmax | SpCas9 | 6 | 6 | 5 | 6 | 6 | 6 | 1 |
| huangcp-CP1012-ABEmax | SpCas9 | 8 | 6 | 6 | 6 | 6 | 6 | 0 |
| huangcp-CP1028-ABEmax | SpCas9 | 8 | 10 | 10 | 10 | 10 | 10 | 0 |
| huangcp-CP1041-ABEmax | SpCas9 | 8 | 7 | 7 | 7 | 7 | 7 | 0 |
| huangcp-CP1249-ABEmax | SpCas9 | 8 | 10 | 10 | 10 | 10 | 10 | 0 |
| tan2019-papap-5 | SpCas9 | 6 | 6 | 6 | 6 | 6 | 6 | 0 |
| tan2019-papapap-7 | SpCas9 | 6 | 5 | 5 | 5 | 5 | 5 | 0 |
| communbiol-dCas12a-A3A | Cas12a | 10 | 4 | 4 | no window | 4 | no window | 0 |
| communbiol-dCas12a-A3A-Y130F | Cas12a | 9 | 4 | 4 | no window | 4 | no window | 0 |
| communbiol-dCas12a-ABE8e | Cas12a | 10 | 4 | 4 | 4 | 4 | 7 | 3 |

**Table S4B.** Buildable candidate count, band edges and per-position candidate density at five sampling seeds.

| Seed | Buildable candidates | Reachable range | pos 1 | pos 2 | pos 3 | pos 4 | pos 5 | pos 6 | pos 7 | pos 8 | pos 9 | pos 10 | pos 11 | pos 12 | pos 13 | pos 14 | pos 15 | pos 16 | pos 17 | pos 18 | pos 19 | pos 20 |
| --- | --- | --- | --- | --- | --- | --- | --- | --- | --- | --- | --- | --- | --- | --- | --- | --- | --- | --- | --- | --- | --- | --- |
| baseline | 185 | 3 to 12 | 0 | 0 | 1 | 1 | 12 | 31 | 13 | 8 | 21 | 70 | 21 | 7 | 0 | 0 | 0 | 0 | 0 | 0 | 0 | 0 |
| seed 1 | 181 | 4 to 12 | 0 | 0 | 0 | 2 | 13 | 30 | 16 | 9 | 26 | 63 | 20 | 2 | 0 | 0 | 0 | 0 | 0 | 0 | 0 | 0 |
| seed 2 | 182 | 4 to 12 | 0 | 0 | 0 | 2 | 11 | 28 | 14 | 8 | 25 | 65 | 24 | 5 | 0 | 0 | 0 | 0 | 0 | 0 | 0 | 0 |
| seed 3 | 182 | 4 to 12 | 0 | 0 | 0 | 1 | 9 | 27 | 12 | 11 | 22 | 80 | 16 | 4 | 0 | 0 | 0 | 0 | 0 | 0 | 0 | 0 |
| seed 4 | 184 | 3 to 11 | 0 | 0 | 1 | 0 | 9 | 28 | 22 | 8 | 22 | 75 | 19 | 0 | 0 | 0 | 0 | 0 | 0 | 0 | 0 | 0 |

**Table S4C.** Predicted mode across the XTEN persistence band, by entry.

| Entry | Observed | 4.5 Å | 5.13 Å | 5.77 Å | 6.4 Å | Spread |
| --- | --- | --- | --- | --- | --- | --- |
| kissling-SpRY-ABE8e-HEK-Plasmid-5d | 6 | 6 | 5 | 5 | 5 | 1 |
| kissling-SpRY-ABE8e-HEK-Plasmid-10d | 6 | 6 | 5 | 5 | 5 | 1 |
| kissling-SpRY-ABE8e-HEK-mRNA-02pmol | 5 | 6 | 5 | 5 | 5 | 1 |
| kissling-SpRY-ABE8e-HEK-mRNA-1pmol | 5 | 6 | 5 | 5 | 5 | 1 |
| kissling-SpRY-ABE8e-HEK-mRNA-5pmol | 5 | 6 | 5 | 5 | 5 | 1 |
| kissling-SpRY-ABE8e-AAVlibrary-AAV-6w | 5 | 6 | 5 | 5 | 5 | 1 |
| kissling-SpRY-ABE8e-AAVlibrary-AAV-12w | 5 | 6 | 5 | 5 | 5 | 1 |
| kissling-SpRY-ABE8e-LentiAAV | 6 | 6 | 5 | 5 | 5 | 1 |
| kissling-SpRY-ABE8e-LentiLNP | 5 | 6 | 5 | 5 | 5 | 1 |
| kissling-SpRY-ABEmax-HEK-Plasmid-5d | 6 | 6 | 5 | 6 | 5 | 1 |
| kissling-SpRY-ABEmax-HEK-Plasmid-10d | 6 | 6 | 5 | 6 | 5 | 1 |
| kissling-SpRY-ABEmax-HEK-mRNA-02pmol | 6 | 6 | 5 | 6 | 5 | 1 |
| kissling-SpRY-ABEmax-HEK-mRNA-1pmol | 6 | 6 | 5 | 6 | 5 | 1 |
| kissling-SpRY-ABEmax-HEK-mRNA-5pmol | 6 | 6 | 5 | 6 | 5 | 1 |
| kissling-SpRY-ABEmax-AAVlibrary-AAV-6w | 6 | 6 | 5 | 6 | 5 | 1 |
| kissling-SpRY-ABEmax-AAVlibrary-AAV-12w | 6 | 6 | 5 | 6 | 5 | 1 |
| kissling-SpRY-ABEmax-LentiAAV | 6 | 6 | 5 | 6 | 5 | 1 |
| kissling-SpRY-ABEmax-LentiLNP | 6 | 6 | 5 | 6 | 5 | 1 |
| kissling-SpG-ABE8e-HEK-Plasmid-5d | 5 | 6 | 5 | 5 | 5 | 1 |
| kissling-SpG-ABE8e-HEK-Plasmid-10d | 5 | 6 | 5 | 5 | 5 | 1 |
| kissling-SpG-ABEmax-HEK-Plasmid-5d | 6 | 6 | 5 | 6 | 5 | 1 |
| kissling-SpG-ABEmax-HEK-Plasmid-10d | 6 | 6 | 5 | 6 | 5 | 1 |
| kissling-SpCas9-ABE8e-HEK-Plasmid-5d | 6 | 6 | 5 | 5 | 5 | 1 |
| kissling-SpCas9-ABE8e-HEK-Plasmid-10d | 6 | 6 | 5 | 5 | 5 | 1 |
| kissling-SpCas9-ABEmax-HEK-Plasmid-5d | 6 | 6 | 5 | 6 | 5 | 1 |
| kissling-SpCas9-ABEmax-HEK-Plasmid-10d | 6 | 6 | 5 | 6 | 5 | 1 |
| hnhx-ABEmax7.10 | 6 | 6 | 5 | 6 | 5 | 1 |
| komor-linker-16 | 6 | 5 | 5 | 5 | 5 | 0 |
| bepigs-BE3 | 6 | 5 | 5 | 5 | 5 | 0 |
| bepigs-pi-16 | 9 | 11 | 11 | 11 | 11 | 0 |
| huangcp-ABEmax | 6 | 6 | 5 | 6 | 5 | 1 |
| huangcp-CP1012-ABEmax | 8 | 6 | 6 | 6 | 6 | 0 |
| huangcp-CP1028-ABEmax | 8 | 10 | 10 | 10 | 10 | 0 |
| huangcp-CP1041-ABEmax | 8 | 7 | 7 | 7 | 7 | 0 |
| huangcp-CP1249-ABEmax | 8 | 10 | 10 | 10 | 10 | 0 |
| communbiol-dCas12a-A3A | 10 | 4 | 4 | 4 |  | 0 |
| communbiol-dCas12a-A3A-Y130F | 9 | 4 | 4 | 4 |  | 0 |
| communbiol-dCas12a-ABE8e | 10 | 4 | 5 | 4 | 4 | 1 |

**Table S5.** Per-entry predicted and measured window position and width, with the signed offset.

| Entry | Ortholog | Anchor site | Linker | Observed mode | Predicted mode | Signed offset | Predicted width | Observed width | Note |
| --- | --- | --- | --- | --- | --- | --- | --- | --- | --- |
| kissling-SpRY-ABE8e-HEK-Plasmid-5d | SpRY | n_terminal | xten-32 | 6 | 5 | -1 | 6 |  |  |
| kissling-SpRY-ABE8e-HEK-Plasmid-10d | SpRY | n_terminal | xten-32 | 6 | 5 | -1 | 6 |  |  |
| kissling-SpRY-ABE8e-HEK-mRNA-02pmol | SpRY | n_terminal | xten-32 | 5 | 5 | 0 | 6 |  |  |
| kissling-SpRY-ABE8e-HEK-mRNA-1pmol | SpRY | n_terminal | xten-32 | 5 | 5 | 0 | 6 |  |  |
| kissling-SpRY-ABE8e-HEK-mRNA-5pmol | SpRY | n_terminal | xten-32 | 5 | 5 | 0 | 6 |  |  |
| kissling-SpRY-ABE8e-AAVlibrary-AAV-6w | SpRY | n_terminal | xten-32 | 5 | 5 | 0 | 6 |  |  |
| kissling-SpRY-ABE8e-AAVlibrary-AAV-12w | SpRY | n_terminal | xten-32 | 5 | 5 | 0 | 6 |  |  |
| kissling-SpRY-ABE8e-LentiAAV | SpRY | n_terminal | xten-32 | 6 | 5 | -1 | 6 |  |  |
| kissling-SpRY-ABE8e-LentiLNP | SpRY | n_terminal | xten-32 | 5 | 5 | 0 | 6 |  |  |
| kissling-SpRY-ABEmax-HEK-Plasmid-5d | SpRY | n_terminal | xten-32 | 6 | 5 | -1 | 4 |  |  |
| kissling-SpRY-ABEmax-HEK-Plasmid-10d | SpRY | n_terminal | xten-32 | 6 | 5 | -1 | 4 |  |  |
| kissling-SpRY-ABEmax-HEK-mRNA-02pmol | SpRY | n_terminal | xten-32 | 6 | 5 | -1 | 4 |  |  |
| kissling-SpRY-ABEmax-HEK-mRNA-1pmol | SpRY | n_terminal | xten-32 | 6 | 5 | -1 | 4 |  |  |
| kissling-SpRY-ABEmax-HEK-mRNA-5pmol | SpRY | n_terminal | xten-32 | 6 | 5 | -1 | 4 |  |  |
| kissling-SpRY-ABEmax-AAVlibrary-AAV-6w | SpRY | n_terminal | xten-32 | 6 | 5 | -1 | 4 |  |  |
| kissling-SpRY-ABEmax-AAVlibrary-AAV-12w | SpRY | n_terminal | xten-32 | 6 | 5 | -1 | 4 |  |  |
| kissling-SpRY-ABEmax-LentiAAV | SpRY | n_terminal | xten-32 | 6 | 5 | -1 | 4 |  |  |
| kissling-SpRY-ABEmax-LentiLNP | SpRY | n_terminal | xten-32 | 6 | 5 | -1 | 4 |  |  |
| kissling-SpG-ABE8e-HEK-Plasmid-5d | SpG | n_terminal | xten-32 | 5 | 5 | 0 | 6 |  |  |
| kissling-SpG-ABE8e-HEK-Plasmid-10d | SpG | n_terminal | xten-32 | 5 | 5 | 0 | 6 |  |  |
| kissling-SpG-ABEmax-HEK-Plasmid-5d | SpG | n_terminal | xten-32 | 6 | 5 | -1 | 4 |  |  |
| kissling-SpG-ABEmax-HEK-Plasmid-10d | SpG | n_terminal | xten-32 | 6 | 5 | -1 | 4 |  |  |
| kissling-SpCas9-ABE8e-HEK-Plasmid-5d | SpCas9 | n_terminal | xten-32 | 6 | 5 | -1 | 6 |  |  |
| kissling-SpCas9-ABE8e-HEK-Plasmid-10d | SpCas9 | n_terminal | xten-32 | 6 | 5 | -1 | 6 |  |  |
| kissling-SpCas9-ABEmax-HEK-Plasmid-5d | SpCas9 | n_terminal | xten-32 | 6 | 5 | -1 | 4 |  |  |
| kissling-SpCas9-ABEmax-HEK-Plasmid-10d | SpCas9 | n_terminal | xten-32 | 6 | 5 | -1 | 4 |  |  |
| hnhx-ABEmax7.10 | SpCas9 | n_terminal | xten-32 | 6 | 5 | -1 | 2 | 5 |  |
| hnhx-ABEmax7.10-dHNH | SpCas9 | n_terminal | xten-32 |  | 6 |  | 4 |  |  |
| hnhx-ABEmax7.10-inserted | SpCas9 | hnh_amino | gs-3 | 10 | 18 | 8 | 4 | 8 |  |
| hnhx-ABE8e-inserted | SpCas9 | hnh_amino | gs-3 |  | 8 |  | 4 |  |  |
| komor-linker-3 | SpCas9 | n_terminal | gs-3 |  | 5 |  | 2 | 3 |  |
| komor-linker-16 | SpCas9 | n_terminal | xten-16 | 6 | 5 | -1 | 5 | 5 |  |
| komor-linker-21 | SpCas9 | n_terminal | gs-21 |  | 5 |  | 4 | 6 |  |
| bepigs-BE3 | SpCas9 | n_terminal | xten-16 | 6 | 5 | -1 | 5 | 5 |  |
| bepigs-pi-16 | SpCas9 | pi_amino | xten-16 | 9 | 11 | 2 | 2 | 11 |  |
| bepigs-pi-8 | SpCas9 | pi_amino | xten-8 |  | 10 |  | 2 | 11 |  |
| bepigs-pi-3 | SpCas9 | pi_amino | xten-3 |  |  |  |  | 11 | every conformation of xten-3 clashes with the editor, over 400000 chains |
| bepigs-ruvc-16 | SpCas9 | ruvc_amino | xten-16 |  | 11 |  | 3 |  |  |
| huangcp-ABEmax | SpCas9 | n_terminal | xten-32 | 6 | 5 | -1 | 2 | 4 |  |
| huangcp-CP1012-ABEmax | SpCas9 | cp1012 | xten-32 | 8 | 6 | -2 | 4 | 9 |  |
| huangcp-CP1028-ABEmax | SpCas9 | cp1028 | xten-32 | 8 | 10 | 2 | 5 | 9 |  |
| huangcp-CP1041-ABEmax | SpCas9 | cp1041 | xten-32 | 8 | 7 | -1 | 5 | 9 |  |
| huangcp-CP1249-ABEmax | SpCas9 | cp1249 | xten-32 | 8 | 10 | 2 | 3 | 9 |  |
| tan2019-xten-16 | SpCas9 | n_terminal | xten-16 |  | 5 |  | 5 | 9 |  |
| tan2019-pap-3 | SpCas9 | n_terminal | polyproline-3 |  | 6 |  | 2 |  |  |
| tan2019-papap-5 | SpCas9 | n_terminal | polyproline-5 | 6 | 6 | 0 | 4 | 3 |  |
| tan2019-papapap-7 | SpCas9 | n_terminal | polyproline-7 | 6 | 5 | -1 | 5 | 2 |  |
| communbiol-dCas12a-A3A | Cas12a | n_terminal | xten-16 | 10 | 4 | -6 | 0 | 10 |  |
| communbiol-dCas12a-A3A-Y130F | Cas12a | n_terminal | xten-16 | 9 | 4 | -5 | 0 | 6 |  |
| communbiol-dCas12a-ABE8e | Cas12a | n_terminal | xten-16 | 10 | 4 | -6 | 3 | 5 |  |
